# Reproductive character displacement explains strengthening of mechanical barriers in damselflies

**DOI:** 10.1101/2023.10.26.564279

**Authors:** Andrea Viviana Ballen-Guapacha, Sandra M. Ospina-Garces, Rosa Ana Sanchez-Guillen

## Abstract

Reinforcing natural selection against maladaptive hybrids can favor the strengthening of premating reproductive isolation driving a pattern of Reproductive Character Displacement (RCD). In a recent study conducted in North-West (NW) (older) Spanish hybrid zone, was detected an asymmetric reinforcement of the mechanical isolation in the reciprocal cross direction between *I. graellsii* males and *I. elegans* females. Furthermore, in the North-Central and Mediterranean (NCM) (younger) hybrid zone was also detected a similar strengthening of the mechanical isolation, consistent with a pattern of asymmetric reinforcement in this hybrid zone as well. In this study, we did geometric morphometrics analyses, shape, and Centroid Size (CS), on male and female secondary sexual traits to investigate whether reinforcement has generated a pattern of RCD of these traits in both hybrid zones. We detected, in the NW hybrid zone, unidirectional RCD (CS) of the male caudal appendages of *I. graellsii*, and bidirectional RCD (shape) of the female prothorax. Consistently with the prediction that the signal of reinforcement may diminish rapidly once reinforcement ceases to operate, a stronger signal of RCD was detected in the NCM than in the NW hybrid region. In this region, was detected unidirectional RCD (CS) of the male caudal appendages which was consistent with the lock-and-key mechanism of genital coevolution, as well as RCD (shape) of the female prothorax of *I. elegans.* Interestingly, our study highlights the importance of using geometric morphometrics to deal with the complexity of reproductive structures and controlling for environmental and geographic factors to investigate RCD.

## INTRODUCTION

Character displacement was defined by Brown and Wilson (1956) as a pattern in which two sympatric species present greater differences in certain ecological or reproductive characters than they do in allopatry. When this pattern is produced by selection to avoid the excessive cost of maladaptive hybridization it is named “Reproductive Character Displacement” (RCD) (Servedio and Noor, 2003; Coyne and Orr, 2004). This process has been earlier named by Dobzhansky (1940) as “reinforcing reproductive isolation”. Available empirical evidence suggests that RCD plays an important role in the evolution of diversity (Schluter, 2000; Pfennig and Pfennig, 2005). This mechanism has been reported in different organisms like insects (Brown and Wilson, 1956; Walker 1974; Kawano 2002), fishes (Schluter, 2000; Crampton et al., 2011; Roth-Monzón et al., 2020), birds (Brown and Wilson, 1956; Diamond et al., 1989; Seddon and Tobias, 2010), reptiles (Melville 2002; Dayan and Simberloff, 2005) and amphibians (Brown and Wilson, 1956; Johanet et al., 2009). Commonly, the presence of RCD in genital divergence between closely related species has been used as support for the lock-and-key hypothesis of genital evolution, i.e., as evidence of natural selection acting on genital morphology to prevent hybridization. Although the number of studies testing RCD on genital divergence has increased and are mainly focused on insects (e.g., Kawano, 2002; 2003; Usami et al., 2006; Augustijnen et al., 2022) and arachnids (Kuntner et al., 2009; Costa-Schmidt and Mellender de Araújo, 2010; Muster and Michalik, 2019), these studies have predominantly focused on male divergence, which have prevented a comprehensive investigation of lock-and-key hypothesis of genital evolution.

In damselflies, mechanical isolation due to the incompatibility to form a tandem position is the strongest component of prezygotic isolation (Sánchez-Guillén et al., 2014a; Nava-Bolaños et al., 2017; Wellenreuther and Sánchez-Guillén 2015; Barnard et al., 2017). In damselflies copula involves two contact points. Firstly, the male grasps the female by her prothorax using his caudal appendages, thus achieving the first contact point known as “tandem position”. The second contact point implies female recognition, through tactile isolation: the female bends her abdomen bringing both genitalia into contact, thus achieving the second contact point known as “wheel position”. There is great variation in the caudal male appendages among odonates. McPeek et al. (2011) observed significant differences in the shape of male cerci among six *Enallagma* species, but not among populations within species. Moreover, McPeek studies in *Enallagma* damselflies (McPeek et al. 2008; 2009) have provided empirical support for “lock-and-key” coevolutionary divergence in damselflies. In their research, McPeek et al. (2008, 2009) identified a correlated evolution between male and female secondary genitalia, specifically, male cerci and female mesostigmal plates. They quantified both tempo and mode of male and female genital evolution and they found a similar pattern of punctuated evolution in both structures, proposing that this pattern might be indicative of a lock-and-key mechanism in the tandem position (see also Paulson, 1974, Masly 2012).

The damselflies *Ischnura elegans* and *I. graellsii* overlap their distributions in north and central Spain, where they are in sympatry (see Figure 1). Currently, *I. elegans* has a patchy distribution in Spain, consistent with a mottled hybrid zone (Sánchez-Guillén et al. 2023). This sympatric distribution comprises two unconnected hybrid zones: the North-West (NW) hybrid zone, which extends along the north-west Spanish coast. *Ischnura elegans* was first discovered in this zone during the early 1980s (Ocharan, 1987). The second hybrid zone, a larger zone known as the North-Central and Mediterranean (NCM) hybrid zone, extends from norther and central Spain and along the Mediterranean coast. The earlier records of *I. elegans* in the Mediterranean zone are before 1900, although certain parts of the central zone, records are more recent, specifically from the early 2000s (Sánchez-Guillén et al., 2021b). In previous genomic studies, on-going hybridization and bilateral introgression were detected in the NW hybrid zone (Sánchez-Guillén et al. 2011; Wellenreuther et al. 2018; Sánchez-Guillén et al. 2023), whereas on-going hybridization but unilateral introgression (thorough *I. graellsii*) was detected in the NCM hybrid region. Recently, Arce-Valdés et al. (2023) tested the reinforcement of reproductive isolation in the NW hybrid zone by comparing the intensity of five prezygotic barriers in crosses between *I. elegans* and *I. graellsii* from allopatry *versus* sympatry, and by measuring the fitness of their hybrids. In the reciprocal cross involving *I. graellsii* males and *I. elegans* females, it was detected that the mechanical barrier implicated in tandem formation was more intense in sympatry than in allopatry, and the hybrids resulting from this cross direction exhibited reduced fitness compared to the parental species. However, in the other reciprocal cross direction, the one between *I. elegans* males and *I. graellsii* females, none of the estimated five prezygotic barriers were more intense in sympatry than in allopatry, and hybrids were not formed in allopatry by the accumulated action of these five reproductive barriers, but were formed in sympatry, possibly due to the purge of BDM incompatibilities. Therefore, Arce-Valdés et al. (2023) detected asymmetric reinforcement occurring only in the reciprocal cross involving *I. graellsii* males and *I. elegans* females. Although the reinforcement has not yet been tested in the NCM hybrid zone, the strength of the mechanical barrier reinforced in the NW hybrid zone was similar in strength to that in the NCM zone (unpublished data). This is consistent with the reinforcement of this mechanical barrier occurring in the NW hybrid zone.

**Figure 1.**
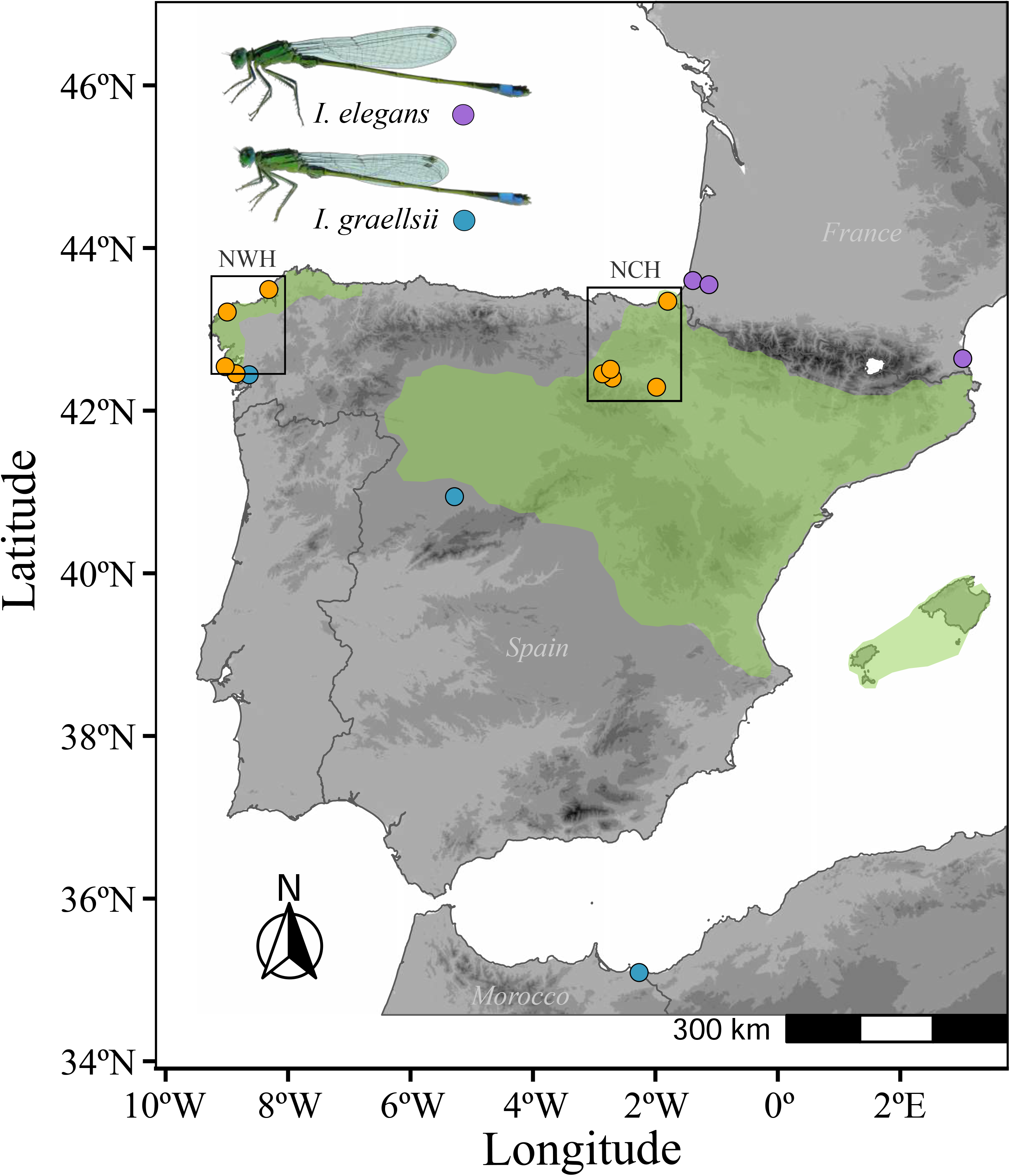
Geographical distribution of *Ischnura elegans* and *I. graellsii* in allopatry and sympatry (contact zone). Yellow dots show sampled localities in both hybrid zones (NW: northwest zone and NCM: north-central and Mediterranean zone), blue dots indicate allopatric *I. graellsii* localities, and purple dots indicate allopatric *I. elegans* localities.

Taking advantage of these features, we investigated whether the reinforcement of mechanical isolation in the reciprocal cross between *I. graellsii* males and *I. elegans* females in the NW and the NCM hybrid zones have generated a pattern of RCD in the secondary-genital divergence of both, *I. graellsii* males and *I. elegans* females. To this end, we performed geometric morphometrics analyses on both shape and Centroid Size (CS) of male and female secondary sexual structures of individuals of *I. elegans* and *I. graellsii* from allopatry and both hybrid zones (NW and NCM). Moreover, because of phenotypic characters may vary along environmental gradients (Goldberg and Lande, 2006), to discriminate between phenotypic differences caused by sympatry from those caused by environmental factors we have controlled for environmental factors (Johanet et al., 2009). Our first prediction is that we will detect a pattern of RCD in *I. graellsii* males and *I. elegans* females within both hybrid zones. This prediction in based on the enhanced mechanical isolation detected in previous studies (Arce-Valdés et al. 2023; unpublished-data) but only in the reciprocal cross between *I. graellsii* males and *I. elegans* females within both hybrid zones. Our second prediction is that RCD could also be detected in *I. graellsii* females and *I. elegans* males within both hybrid zones. This prediction is based on the lock-and-key coevolution of the genitalia, which will drive a genital coevolution as a mechanism to maintain the integrity of species boundaries and to prevent hybridization (see Sloan and Simmons, 2019). Our third prediction is that RCD will be unilateral or asymmetric between species. This prediction is based on that the strength of RCD depends on the interplay of several non-mutually exclusive factors: 1) the symmetry/asymmetry of reinforcement of the reproductive isolation (cf. Cooley, 2007; Yikweon, 2008); 2) proximity to parental populations, since hybrid populations closer to parental populations experience higher levels of gene flow (Liou and Price 1994; Pfennig and Ryan 2006); the 3) the abundance of both parental species, since rarer species have a greater chance of encountering/mating with the common species than *vice versa* (Peterson et al., 2005; Hochkirch et al., 2007). Our fourth prediction is that the signal of RCD will be stronger in the NCM than in the NW hybrid region because the signal of reinforcement may diminish rapidly once reinforcement ceases to operate. This prediction is based on higher levels of interspecific gene flow detected in the NW compared to NCM hybrid region (see Sánchez-Guillén et al. 2023). Our fifth prediction is that RCD will also vary between populations because of differences in local species proportions (see Sánchez-Guillén et al. 2023).

## METHODOLOGY

### Population samplings

During summers from 2003 to 2021, adult males and females were collected from 15 localities belonging to six allopatric localities, four localities from the NW hybrid zone, and five localities from the NCM hybrid zone (Table 1; Figure 1). Populations from both hybrid zones were selected for this study based on our knowledge of reproductive isolation and hybridization patterns from previous studies. Arce-Valdes et al. (2023) detected the strengthening of the mechanical isolation due to the mismatch between male and female secondary genital structures to form the tandem in both conspecific crosses of both species, and heterospecific crosses in sympatry with respect to allopatry. From each locality, we sampled between 14-31 adult males and 9-28 adult females of both species depending on the dominant species in each locality (Table 1). Sampled specimens were preserved in ethanol (96%) and stored at −20°C.

**Table 1.** Sample details of *I. elegans* and *I. graellsii* for the morphometric analysis. Species proportions indicate species proportion in each sampled population. N males and N females indicated the number of samples included in the morphometric analyses. The distribution indicates allopatric and sympatric populations: NW (North-west) and NCM (North-central and Mediterranean) hybrid zones.

| Species | Locality | Distribution | Species proportion | Males | Females | Year | Latitude | Longitude |
| --- | --- | --- | --- | --- | --- | --- | --- | --- |
| *I. elegans* | Marais D'Orx, France | Allopatry | 100% | 25 | 7 | 2015 | 43°35'58"N | 1°23'30"W |
| *I. elegans* | Sablière, France | Allopatry | 100% | 26 | 8 | 2016 | 46°49'53"N | 1°51'53"W |
| *I. elegans* | St. Cyprien, France | Allopatry | 100% | 26 | 25 | 2016 | 42°37'07"N | 3°00'16"W |
| *I. graellsii* | Gamillazo, Salamanca | Allopatry | 100% | 14 | 20 | 2021 | 40°57'22"N | 5°17'21"W |
| *I. graellsii* | Saïdia, Morocco | Allopatry | 100% | - | 16 | 2009 | 35°05'27"N | 2°15'57"W |
| *I. graellsii* | Alba, Galicia | Allopatry | 100% | 16 | 9 | 2003 | 42°26'30"N | 8°38'05"W |
| *I. elegans* | Doniños, Galicia | Sympatry (NW) | 100% | 24 | 23 | 2016 | 43°29'27"N | 8°18'29"W |
| *I. elegans* | Laxe, Galicia | Sympatry (NW) | 99-100% | 27 | 22 | 2018 | 43°12'44"N | 8°59'43"W |
| *I. graellsii* | Cachadas, Galicia | Sympatry (NW) | 99-100% | 31 | 28 | 2018-2019 | 42°27'12"N | 8°50'52"W |
| *I. graellsii* | Xuño, Galicia | Sympatry (NW) | 90-100% | 30 | 13 | 2005, 2013, 2016 | 42°37'59"N | 9°02'19"W |
| *I. elegans* | Perdiguero, Rioja | Sympatry (NCM) | 71-90% | 19 | 20 | 2018 | 42°17'06"N | 1°58'55"W |
| *I. elegans* | Plaiaundi, Rioja | Sympatry (NCM) | 99-100% | 16 | - | 2016 | 43°20'44"N | 1°47'40"W |
| *I. elegans* | Villar, Rioja | Sympatry (NCM) | 74-100% | 25 | 22 | 2018 | 42°24'03"N | 2°42'15"W |
| *I. graellsii* | Valpierre, Rioja | Sympatry (NCM) | 95-100% | 30 | 22 | 2018 | 42°27'11"N | 2°51'48"W |
| *I. graellsii* | Mateo, Rioja | Sympatry (NCM) | 95-100% | 21 | 33 | 2018 | 42°30'34"N | 2°44'05"W |

### Geometric morphometrics analyses

We used a two-dimensional geometric morphometric approach to investigate the RCD of genital structures because it allows to break down the complexity in shape and CS of the studied reproductive structures (Klingenberg, 2016). Thus, our analyses investigated RCD in two views (lateral and posterior) of both shape and CS of male caudal appendages and female prothorax because these are structures that can influence the ability to achieve copulation and contribute to defining species-specificity (Simmons and García-Gonzalez, 2011; Song, 2009). In fact, morphological differences in the male’s caudal appendages and the female’s prothorax are used in morphological identification of these species (Askew, 1988; Monetti et al., 2002). In addition, both structures are involved in the first point of contact during damselfly reproduction which is known as the tandem position. During the tandem male grasps the female by her thorax by using his caudal appendages to achieve this position (Supplementary Figure S1A-B), thereafter, the second point of contact implies a prior recognition of the female, which bends her abdomen putting into contact with both genitalia, thus achieving the copula.

Photographs of the posterior and lateral view of the male’s caudal appendages and the female’s prothorax were obtained under a stereoscopic microscope (Zeiss Stemi 305) with an integrated camera (Axiocam ERc5s). The posterior view of the male’s caudal appendages shape was sampled with a combination of eight landmarks and 80 semi-landmarks in four curves (left and right sides, Supplementary Figure S2A-B), which were assigned above and below each of the cerci. The lateral view was sampled with seven landmarks and 36 semi-landmarks in two curves which were assigned on the cercus and the paraproct of the left side (Supplementary Figure S2C-D). Likewise, the posterior view of the female’s prothorax shape was sampled with a combination of four landmarks and 21 semi-landmarks in one curve (Supplementary Figure S3A-B), which was assigned surrounding the pronotum. The left lateral view was sampled with 4 landmarks and 19 semi-landmarks in one curve (Supplementary Figure S3C-D), also surrounding the pronotum Cartesian coordinates of each posterior and lateral view of males and females were registered with tpsDIG2 version 2.04 (Rohlf, 2006): 88 landmarks in the posterior view of the caudal appendages, and 43 landmarks in lateral view; 25 landmarks in posterior view of pronotum and 23 landmarks in lateral view; in which x and y coordinates of the geometric configurations were obtained for each image. The configuration of the landmarks on each of the male and female reproductive structures of both species is described in table S1.

From these, a Generalized Procrustes Analysis (GPA; Rohlf and Slice, 1990) was generated to superimpose landmark configurations and variation due to differences in translation, orientation, and size was removed followed by a thin-plate spline analysis (Bookstein 1991) using the geomorph package version 4.0.4 (Adams et al. 2022). The GPA translates all samples to an origin, rescales them to a CS, and rotates them by least squares to the coordinates of the corresponding points, aligning them as closely as possible (Adams et al., 2017). Semilandmarks were aligned using the Bending Energy minimization criteria, which optimize the semilandmarks position with the lowest deformation energy from a consensus curve of reference (Gunz and Mitteroecker 2013). Additionally, shape variation of both reproductive structures (male’s caudal appendages and female’s prothorax) was then evaluated graphically along the first two axes of a Principal Components Analysis (PCA) and through deformation grids to facilitate anatomical description of any implied morphological changes using the same geomorph package version 4.0.4 (Adams et al. 2022).

### Morphological variation: environmental and geographic factors and/or reinforcement of RI?

Ecological changes in habitat or resource use may generate changes in the species that promote RCD (Pfennig and Pfennig, 2009). Earlier studies in odonates, for instance, have shown that phenotypic variables, such as body size or wing size and shape show a latitudinal variation (Johansson, 2003; Hassall et al., 2008). Since our allopatric and sympatric studied populations are distributed in a latitudinal and longitudinal gradient, environmental factors may also be responsible for explaining their phenotypic characters (Goldberg and Lande, 2006). Thus, to discriminate between phenotypic differences caused by sympatry from those caused by environmental/geographic we tested whether environmental and geographic factors and sympatry might be correlated with the morphological characters measured.

To this end, we analyzed five environmental (precipitation, annual mean temperature, maximum temperature, minimum temperature, and elevation) and two geographic (latitude and longitude) factors. Environmental variables were chosen from those identified in previous studies as variables that contribute to the geographical distribution of *I. elegans* and that are involved in odonate selection processes (Lancaster et al., 2015; Dudaniec et al., 2018; 2021; Wellenreuther et al., 2012). Climate data were generated using bioclimatic parameters from the Worldclim database version 1.4 (Fick and Hijmans, 2017) (at 1km cell resolution). Rasters from each variable were obtained using the function ‘getData’ from the ‘raster’ package. Furthermore, we explored a correlation between environmental variables using the ‘PerformanceAnalytics’ package version 2.0.4 (Peterson et al., 2022). Thus, for both environmental and geographic data, when two variables presented an absolute value of correlation coefficients (|r|)>0.7, one of them was not included in the same model because the relationship between these is considered strong (see Dormann et al., 2013).

Finally, we fitted separate gaussian generalized linear models (GLMs) with an identity link function for the shape (described by the first principal component of shape variance to each geometric configuration), and for the CS of male and female reproductive structures in each species. All models included the geographic variables (latitude and longitude) because in all data groups, they showed correlation coefficient values lower than 0.7 (Table S2), then we added those environmental variables with less than 0.7 absolute correlation coefficient with both geographic variables. When two environmental variables had low correlation coefficients with the geographic variables but were highly correlated between them, we selected the variable with the overall lowest correlation coefficients with the geographic variables. For males and females of *I. elegans* and for males of *I. graellsii*, the explanatory variables of the GLMs included one environmental variable (annual mean temperature), the two geographical variables (latitude and longitude), and the study zones (allopatry, NW and NCM hybrid zones) as a fixed effect to test for zone variation in individual phenotypic responses to these environmental factors. While for *I. graellsii* females, the explanatory variables used in the model were the elevation, the two geographical variables, and the study zones (allopatry, NW and NCM hybrid zones) as a fixed effect. During model selection, Corrected Akaike Information Criterion (AICc) and the significance of fixed effects were used to distinguish among competing models. All analyses were conducted using the statistical software R-project (version 3.6.2, R Core Team, 2019).

### Reproductive character displacement by regions

To tested our first four predictions: 1) RCD will be detected in *I. graellsii* males and *I. elegans* females by reinforcement of RI; 2) RCD could also be detected in *I. graellsii* females and *I. elegans* females by lock-and-key coevolution; 3) RCD could be unilateral or asymmetric between species because it depends on the interplay of several non-mutually exclusive factors; and 4) RCD will be more pronounced in the younger NCM hybrid zone because of the signal of reinforcement may diminish rapidly once reinforcement ceases to operate, in the following manner.

Because RCD is a pattern whereby reproductive characters can already differ or not in allopatry (Slatkin, 1980; Liou and Price, 1994), but the variance in distance between these reproductive characters should be larger in sympatry than in allopatry, first we measured the distance variance between *I. elegans* and *I. graellsii* from allopatry and from each hybrid zone. When the variance in distance was larger in the hybrid zone compared to the allopatric zone (distance in sympatry - distance in allopatry 0) this provided evidence of RCD (see Table 2). Second, in cases where RCD was detected given its potential for being unilateral or bilateral, occurring in either one or both species we conducted intraspecific comparisons (distance in sympatry *versus* distance in allopatry). Therefore, when the variance in distance was significantly different only in one intraspecific comparison, we inferred that RCD was asymmetric (occurring in one species), and when the variance in distance was significantly different in both intraspecific comparisons, we inferred that RCD was symmetric.

**Table 2.**
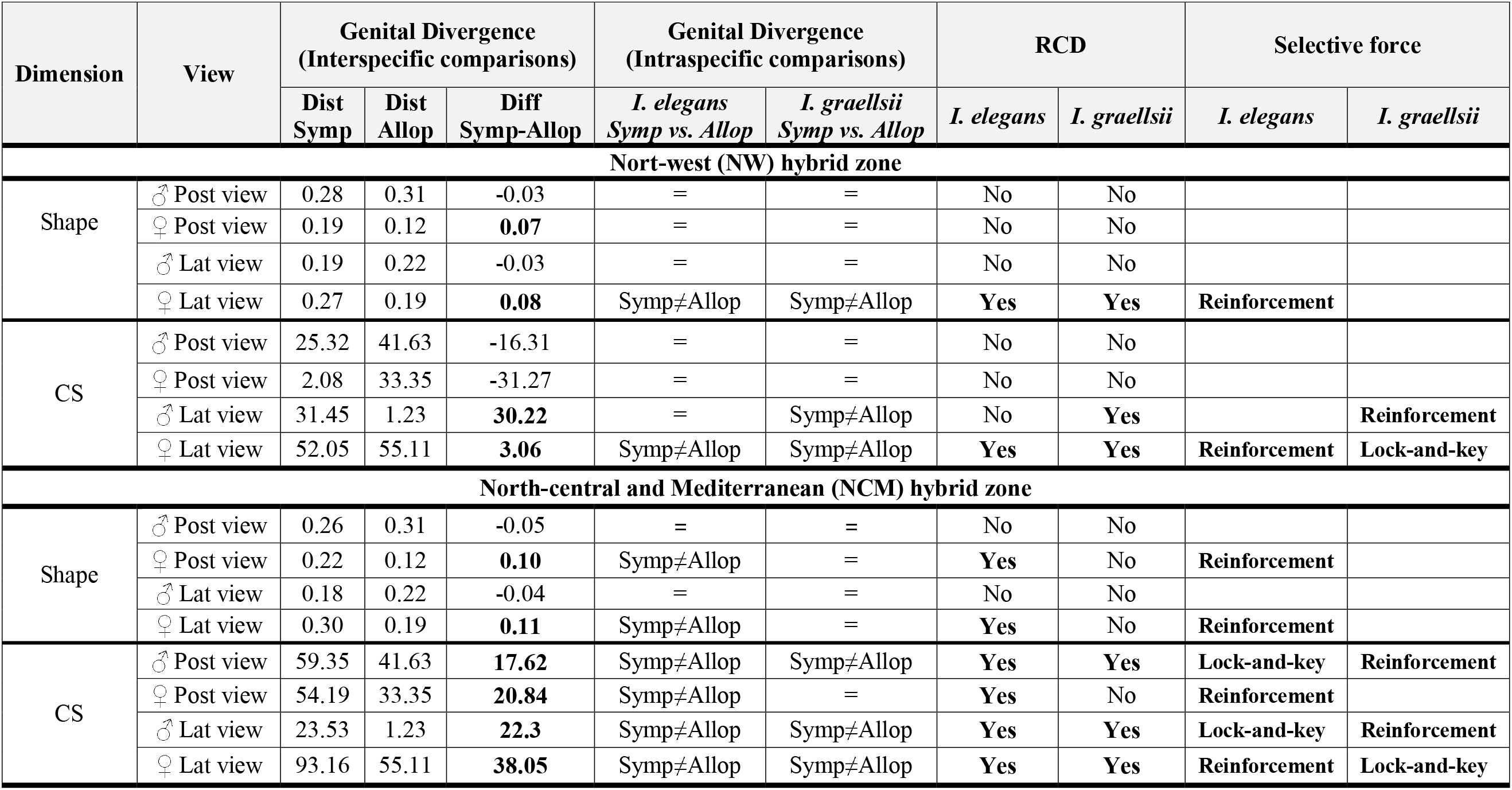
Reproductive character displacement (RCD) in *I. elegans* and *I. graellsii* from both hybrid zones. First and second columns indicate whether analyses correspond to the shape or CS, sexes (male or females) and views (lateral or posterior). Third and fourth columns include interspecific and intraspecific comparisons. Fifth column indicate when genital variance was consistent with a pattern of RCD in one or both species. Sixth column indicates the selective force explaining the RCD pattern.

| Dimension | View | Genital Divergence<br>(Interspecific comparisons) |  |  | Genital Divergence<br>(Intraspecific comparisons) |  | RCD |  | Selective force |  |
| --- | --- | --- | --- | --- | --- | --- | --- | --- | --- | --- |
|  |  | Dist<br>Symp | Dist<br>Allop | Diff<br>Symp-Allop | *I. elegans*<br>*Symp* vs. *Allop* | *I. graellsii*<br>*Symp* vs. *Allop* | *I. elegans* | *I. graellsii* | *I. elegans* | *I. graellsii* |
| Nort-west (NW) hybrid zone |  |  |  |  |  |  |  |  |  |  |
| Shape | ♂ Post view | 0.28 | 0.31 | -0.03 | = | = | No | No |  |  |
|  | ♀ Post view | 0.19 | 0.12 | **0.07** | = | = | No | No |  |  |
|  | ♂ Lat view | 0.19 | 0.22 | -0.03 | = | = | No | No |  |  |
|  | ♀ Lat view | 0.27 | 0.19 | **0.08** | Symp≠Allop | Symp≠Allop | **Yes** | **Yes** | **Reinforcement** |  |
| CS | ♂ Post view | 25.32 | 41.63 | -16.31 | = | = | No | No |  |  |
|  | ♀ Post view | 2.08 | 33.35 | -31.27 | = | = | No | No |  |  |
|  | ♂ Lat view | 31.45 | 1.23 | **30.22** | = | Symp≠Allop | No | **Yes** |  | **Reinforcement** |
|  | ♀ Lat view | 52.05 | 55.11 | **3.06** | Symp≠Allop | Symp≠Allop | **Yes** | **Yes** | **Reinforcement** | **Lock-and-key** |
| North-central and Mediterranean (NCM) hybrid zone |  |  |  |  |  |  |  |  |  |  |
| Shape | ♂ Post view | 0.26 | 0.31 | -0.05 | = | = | No | No |  |  |
|  | ♀ Post view | 0.22 | 0.12 | **0.10** | Symp≠Allop | = | **Yes** | No | **Reinforcement** |  |
|  | ♂ Lat view | 0.18 | 0.22 | -0.04 | = | = | No | No |  |  |
|  | ♀ Lat view | 0.30 | 0.19 | **0.11** | Symp≠Allop | = | **Yes** | No | **Reinforcement** |  |
| CS | ♂ Post view | 59.35 | 41.63 | **17.62** | Symp≠Allop | Symp≠Allop | **Yes** | **Yes** | **Lock-and-key** | **Reinforcement** |
|  | ♀ Post view | 54.19 | 33.35 | **20.84** | Symp≠Allop | = | **Yes** | No | **Reinforcement** |  |
|  | ♂ Lat view | 23.53 | 1.23 | **22.3** | Symp≠Allop | Symp≠Allop | **Yes** | **Yes** | **Lock-and-key** | **Reinforcement** |
|  | ♀ Lat view | 93.16 | 55.11 | **38.05** | Symp≠Allop | Symp≠Allop | **Yes** | **Yes** | **Reinforcement** | **Lock-and-key** |

Statistical analyses were done in the following manner. Firstly, to assess the statistical hypothesis investigating RCD patterns of shape variation and covariation for the set of (Procrustes-aligned) coordinates, explained by CS, geographic group (allopatry, NW and NCM hybrid zones) and their interaction we performed a procrustes ANOVA model using the ‘procD.lm’ function from the geomorph package version 4.0.4 (Adams et al., 2022), which calculates the Procrustes distances variance explained by each factor of the model. Secondly, when the group factor showed statistical significance, we complemented this analysis with a pairwise evaluation using the ‘permudist’ function from the ‘morpho’ package version 2.8 (Schlager, 2017). This function compares the distance between two group means with the distances obtained by random assignment (here 1,000 permutations) of observations to the compared groups (Schlager, 2017). Thirdly, to assess differences in CS we used the function ‘lm.rrpp’ from the RRPP package (Collyer and Adams et al., 2018; 2021) to perform the linear models. When the group factor showed statistical significance, we conducted posthoc pairwise comparisons to detect which groups were statistically different using the ‘pairwise’ function.

### Reproductive character displacement by localities

To test our fifth prediction, that RCD will vary between populations because of differences in local species proportions (see Sánchez-Guillén et al. 2023) we separately analyzed the shape and CS of male caudal appendages and female prothorax in each population.

To assess the statistical hypothesis investigating RCD patterns of shape variation and covariation for the set of (Procrustes-aligned) coordinates, explained by CS, locality, and their interaction we performed a procrustes ANOVA model using the ‘procD.lm’ function from the geomorph package version 4.0.4 (Adams et al., 2022). When the locality showed statistical significance, we complemented this analysis with a pairwise evaluation using the ‘permudist’ function from the ‘morpho’ package version 2.8 (Schlager, 2017). To assess the statistical hypothesis investigating RCD patterns of CS variation we used the function ‘lm.rrpp’ from the RRPP package (Collyer and Adams et al., 2018; 2021) to perform the linear models. When the locality factor showed statistical significance, we conducted posthoc pairwise comparisons to detect which localities were statistically different using the ‘pairwise’ function.

### Lock-and key coevolution of male-female genitalia

We evaluated the role of the lock-and-key on the coevolution of male and female genitalia of *I. elegans* and *I. graellsii* males and females given RCD was detected in both males and females (see Table 2 for detailed results). To this end, we followed two approximations.

Our first approximation was based on that the lock-and-key mechanism, which predicts that genital coevolution results from reinforcement selection, as the formation of new species requires reproductive isolation (Masly 2012). Reinforcing natural selection represents a stabilizing force that acts to strengthen RI between species. As a result, it is expected to promote allometric relationships between body size and genital size. This occurs because it pushes to maintain an equilibrium between these two traits, ensuring optimal reproductive compatibility. To test this prediction, we examined a subset of the data, which included one allopatric population of each species (Cyprien for *I. elegans* and Gamillazo for *I. graellsii*) and two populations from each hybrid zone (NW hybrid zone: Laxe for *I. elegans* and Cachadas for *I. graellsii*, and the NCM hybrid zone: Perdiguero for *I. elegans* and Valpierre for *I. graellsii*). We investigated whether male and female genital size changed allometrically in relation to body length by using wing length as a proxy for body size (Nava et al., 2011; 2014; Córdoba et al., 2015). This approach was necessary due to abdominal deformation during preservation in ethanol. We included a set of 10 males and 10 females from each population, zone, and species. The wings were dissected using fine scissors and examined under a stereoscopic microscope (Zeiss Stemi 305), and we measured the hindwing length between two points: the arculus and the proximal corner of the pterostigma (Carchini et al., 2000) using IMAGEJ software (version 1.53k, National Institutes of Health, MD, USA). To keep a constant measurement error, the wing length was measured by the same person. We performed a Pearson correlation between CS genital size and hindwing length. Both the CS data and the wing length data were log-transformed to linearize the relationship between genital traits and body size indicators. We used R software (R Development Core Team 2009) for statistical analysis.

Secondly, we directly examined the lock-and-key genital coevolution (shape and CS, and both views) of the complementary male caudal appendages and female prothorax in allopatry as well as in both hybrid zones. To test this prediction, we examined a subset of data, in which we select the same number of males as females. Using randomization in R, we adjusted the number of pairs according to the minimum number of individuals of either sex in each population. To assess the genital coevolution of the shape of male and female genital structures, we conducted a two-block partial least squares analysis on Procrustes shape variables using paired samples with 100 iterations, utilizing the ‘two.b.pls’ function from the geomorph package version 4.0.4 (Adams et al. 2022). To assess the genital coevolution of the CS of male and female genital structures, we performed a Pearson correlation with 100 iterations. All analyses were done with the statistical software R-project (version 3.6.2, R Core Team, 2019).

## RESULTS

### Morphological variation: environmental and geographic factors and/or reinforcement of RI?

A total of 192 *I. elegans* males and 137 *I. graellsii* males were included in the morphometric analyses of male caudal appendages, and a total of 134 *I. elegans* females and 135 *I. graellsii* females were included in the morphometric analyses of the prothorax. Table 3 includes all statistical GLM results explaining which part of the morphological variation detected on male and female secondary genitalia was explained by the annual mean temperature or elevation (as they were the only uncorrelated and statistically significant environmental factors), the two geographic factors (latitude and longitude) and the distribution type (allopatry, NW and NCM hybrid zones).

**Table 3.** Influence of environmental and geographical factors on the male and female reproductive characters of *Ischnura elegans* and *I. graellsii* damselflies from allopatric and hybrid zones. Estimated regression parameters, estándar errors, t-values and P-values for the Gaussian GLMs applied on the shape and the CS of reproductive structures.

| Morphological character | Character view |  | Environmental/geographical factors | *I. elegans* |  |  |  | *I. graellsii* |  |  |  |
| --- | --- | --- | --- | --- | --- | --- | --- | --- | --- | --- | --- |
|  |  |  |  | Estimate | Std. Error | t value | P-value | Estimate | Std. Error | t value | P-value |
| Caudal appendage | Posterior | CS | Annual mean temperature | -28.018 | 7.780 | -3.601 | **<0.001** | NS | NS | NS | NS |
|  |  | PC1 | Annual mean temperature | -0.018 | 0.006 | -2.958 | **0.003** | NS | NS | NS | NS |
|  |  | CS | Longitude | -8.715 | 3.568 | -2.443 | **0.015** | 9.674 | 4.797 | 2.017 | **0.045** |
|  |  | PC1 | Longitude | 0.007 | 0.003 | 2.479 | **0.014** | NS | NS | NS | NS |
|  |  | CS | NW Zone | -94.936 | 31.718 | -2.993 | **0.003** | NS | NS | NS | NS |
|  |  | PC1 | NW Zone | 0.081 | 0.025 | 3.297 | **0.001** | 0.039 | 0.011 | 3.623 | **<0.001** |
|  |  | PC1 | NCM Zone | NS | NS | NS | NS | 0.051 | 0.011 | 4.638 | **<0.001** |
|  |  | PC1 | Allopatry Zone | -0.028 | 0.010 | -2.903 | **0.004** | NS | NS | NS | NS |
| Caudal appendage | Lateral | PC1 | Annual mean temperature | -0.025 | 0.011 | -2.317 | **0.021** | NS | NS | NS | NS |
|  |  | CS | Longitude | -11.331 | 3.216 | -3.523 | **<0.001** | 18.650 | 8.255 | 2.259 | **0.025** |
|  |  | PC1 | Longitude | 0.011 | 0.005 | 2.141 | **0.033** | NS | NS | NS | NS |
|  |  | CS | NW Zone | -119.91 | 28.588 | -4.194 | **<0.001** | 41.805 | 12.375 | 3.378 | **<0.001** |
|  |  | PC1 | NW Zone | 0.129 | 0.043 | 2.988 | **0.003** | NS | NS | NS | NS |
| Prothorax | Posterior | CS | Latitude | NS | NS | NS | NS | 3.388 | 1.58 | 2.144 | **0.033** |
|  |  | CS | Longitude | -13.664 | 5.216 | -2.62 | **0.009** | NS | NS | NS | NS |
|  |  | PC1 | Longitude | NS | NS | NS | NS | -0.003 | 0.001 | -2.405 | **0.017** |
|  |  | CS | NW Zone | -149.918 | 45.423 | -3.3 | **0.001** | NS | NS | NS | NS |
|  |  | PC1 | NCM Zone | -0.058 | 0.019 | -3.142 | **0.002** | NS | NS | NS | NS |
|  |  | PC1 | Allopatry Zone | 0.077 | 0.014 | 5.448 | **<0.001** | NS | NS | NS | NS |
| Prothorax | Lateral | PC1 | Latitude | NS | NS | NS | NS | 0.009869 | 0.002746 | 3.593 | **<0.001** |
|  |  | CS | Longitude | -4.148 | 1.869 | -2.219 | **0.028** | NS | NS | NS | NS |
|  |  | PC1 | Longitude | NS | NS | NS | NS | 0.005 | 0.002 | 2.263 | **0.025** |
|  |  | CS | NW Zone | NS | NS | NS | NS | 10.454 | 3.944 | 2.651 | **0.009** |
|  |  | CS | NCM Zone | 36.018 | 8.897 | 4.048 | **<0.001** | 13.116 | 3.771 | 3.478 | **<0.001** |
|  |  | PC1 | NCM Zone | 0.062 | 0.021 | 2.989 | **0.003** | *NS* | *NS* | *NS* | *NS* |
|  |  | PC1 | Allopatry Zone | -0.136 | 0.016 | -8.573 | **<0.001** | *NS* | *NS* | *NS* | *NS* |

In *I. elegans* males GLM modeling and scoring using the AICc suggested that the morphological variation of the shape, at both views, of their caudal appendages was influenced by the annual mean temperature, the longitude, and the distribution type, while the morphological variation of the CS differed between views: the lateral view was influenced by the annual mean temperature, the longitude, and the distribution type; while for the lateral view only the longitude and the distribution type were included in the best model. According to the post-hoc comparisons, based on the shape (both views) the allopatric zone was different from both hybrid zones (Table 3 and S3), while based on the CS (both views), the allopatric zone was only different from the NCM hybrid zone (Table 3 y S3). Otherwise, in *I. elegans* females GLM modeling and scoring using the AIC suggested that the morphological variation of the shape, at both views, of their prothorax was not influenced by any environmental or geographical variable, only the distribution type was included in the best model, while the morphological variation of the CS, at both views, was influenced by the longitude and the distribution type. Similarly, to that detected in males, post-hoc comparisons based on the shape (both views) the allopatric zone was different from the hybrid zones (Table 3 y S3), while based on the CS (both views), the allopatric zone was only different from the NCM hybrid zone (Table 3 y S3).

In *I. graellsii* males GLM modeling and scoring using the AICc suggested that the morphological variation of the lateral view of the shape of their caudal appendages was influenced by the distribution type, while the morphological variation of the CS differed between views: the lateral view was influenced by the longitude and the distribution type, while for the posterior view only the distribution type was included in the best model. According to the post-hoc comparisons, based on the shape (lateral view) the allopatric zone was different from both hybrid zones (Table 3 and S3), while based on the CS the allopatric zone was only different from the NCM hybrid zone for the posterior view, and was only different from the NW hybrid zone for the lateral view (Table 3 y S3). Otherwise, in *I. graellsii* females GLM modeling and scoring using the AIC suggested that the morphological variation of the shape, at both views, of their prothorax was not influenced by any environmental variable or the distribution type, only by the geographical variables were included in the best model: the longitude at both views and the latitude only the lateral view (Table 3), similarly, the morphological variation of the CS differed between views: for the posterior view, the best model included only the latitude, while for the lateral view, the best model included only the distribution type. Post-hoc comparisons based on the CS (lateral vie) indicated differences between the allopatric zone and the hybrid zones (Table 3 y S3).

### Shape variation of male caudal appendages

PCA plots of the two views (posterior and lateral) showed a clear separation between species in the three zones (Table S4, Figure 2A-F). Procrustes ANOVA showed statistically significant differences between *I. elegans* and *I. graellsii*: 1) from the allopatric zone (posterior view: F=326.66, *p*<0.001, d=0.31; lateral view: F=144.25, *p*<0.001, d=0.22); 2) from the NCM hybrid zone (posterior view: F=226.17, *p*<0.001, d=0.26; lateral view: F=90.29, *p*<0.001, d=0.18); and 3) from the NW hybrid zone (posterior view: F=318.78, *p*<0.001, d=0.28; lateral view: F=89.06, *p*<0.001, d=0.19) (Table S5-S6). However, distance variance (both views) did not support the RCD, as the distance variance in both hybrid zones the NW and the NCM hybrid zones was not larger than that in the allopatric zone (distance sympatry - distance allopatry 0) (Table 2).

**Figure 2.**
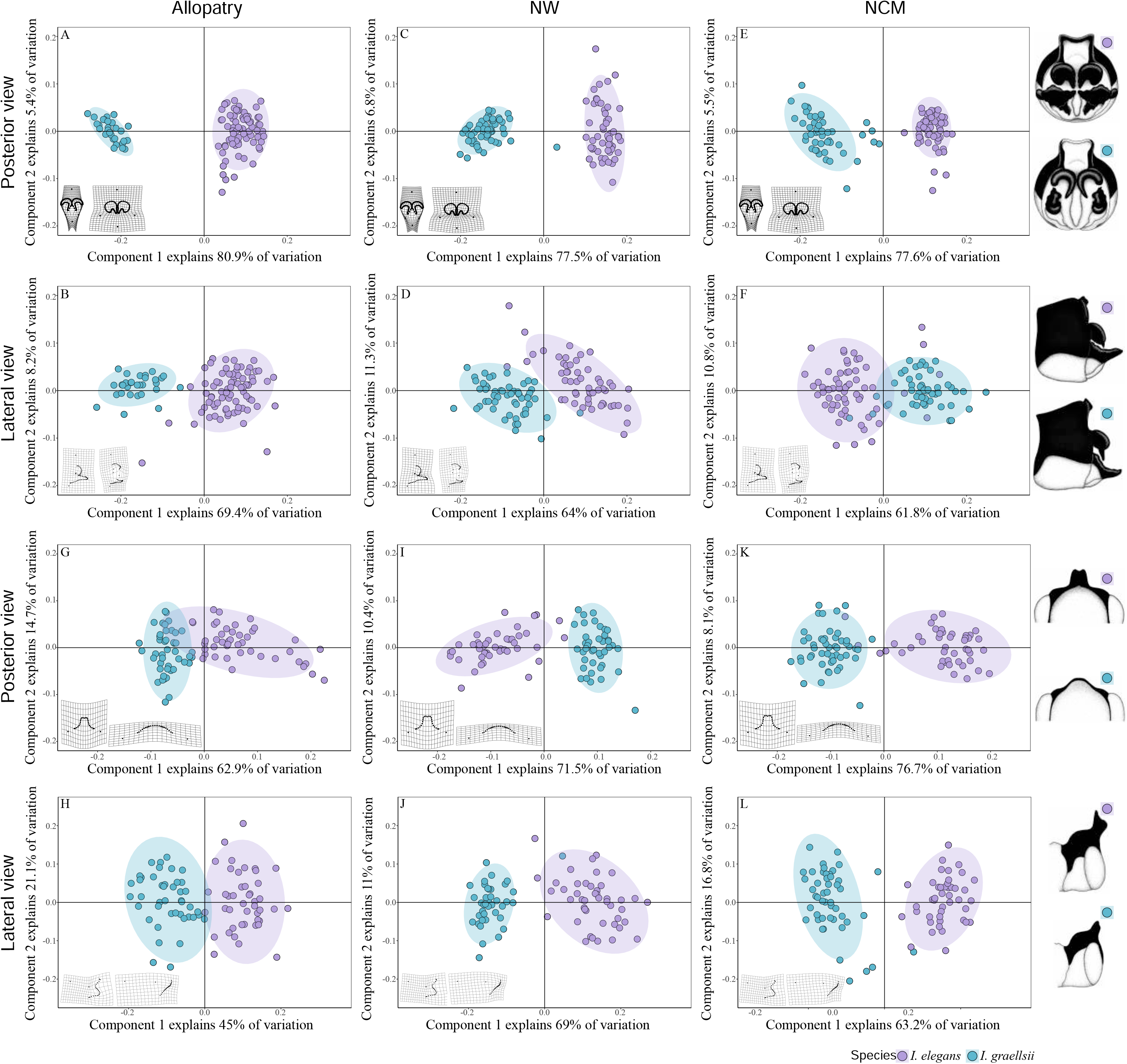
PCA plots showing the variation of males’ caudal appendage in both species (*Ischnura elegans* and *I. graellsii*) in the allopatry regions (A, posterior view and D, lateral view), in the northwest hybrid region (B, posterior view and E, lateral view) and the north-central region (C, posterior view and F, lateral view), and of female’s prothorax (G, posterior view and J, lateral view), in the northwest hybrid region (H, posterior view and K, lateral view) and the north-central region (I, posterior view and L, lateral view).

#### CS variation of male caudal appendages

Procrustes ANOVA showed statistically significant differences between *I. elegans* and *I. graellsii* from allopatry, particularly in the posterior view (posterior view F=18.603, *p*<0.001, d=41.634; lateral view F=0.017, *p*=0.894, d=1.231), and from the NCM (posterior view: F=29.581, *p*<0.001, d=59.347; lateral view: F=5.151, *p*=0.018, d=23.529) and NW (posterior view: F=6.465, *p*=0.013, d=25.324; lateral view: F=13.629, *p*<0.001, d=31.454) hybrid zones (Tables S7-S8, Figure 3A-B). The CS had significant but low effects in all the models (6 to 16% of shape variance, *p*<0.01) and the lowest variances were explained by the interaction between CS and groups (2 to 4%). Distance variance analysis confirmed RCD in both views in the NCM but only in the lateral view in the NW hybrid zone, where the difference between sympatry and allopatry distances was greater than zero (Table 2).

**Figure 3.**
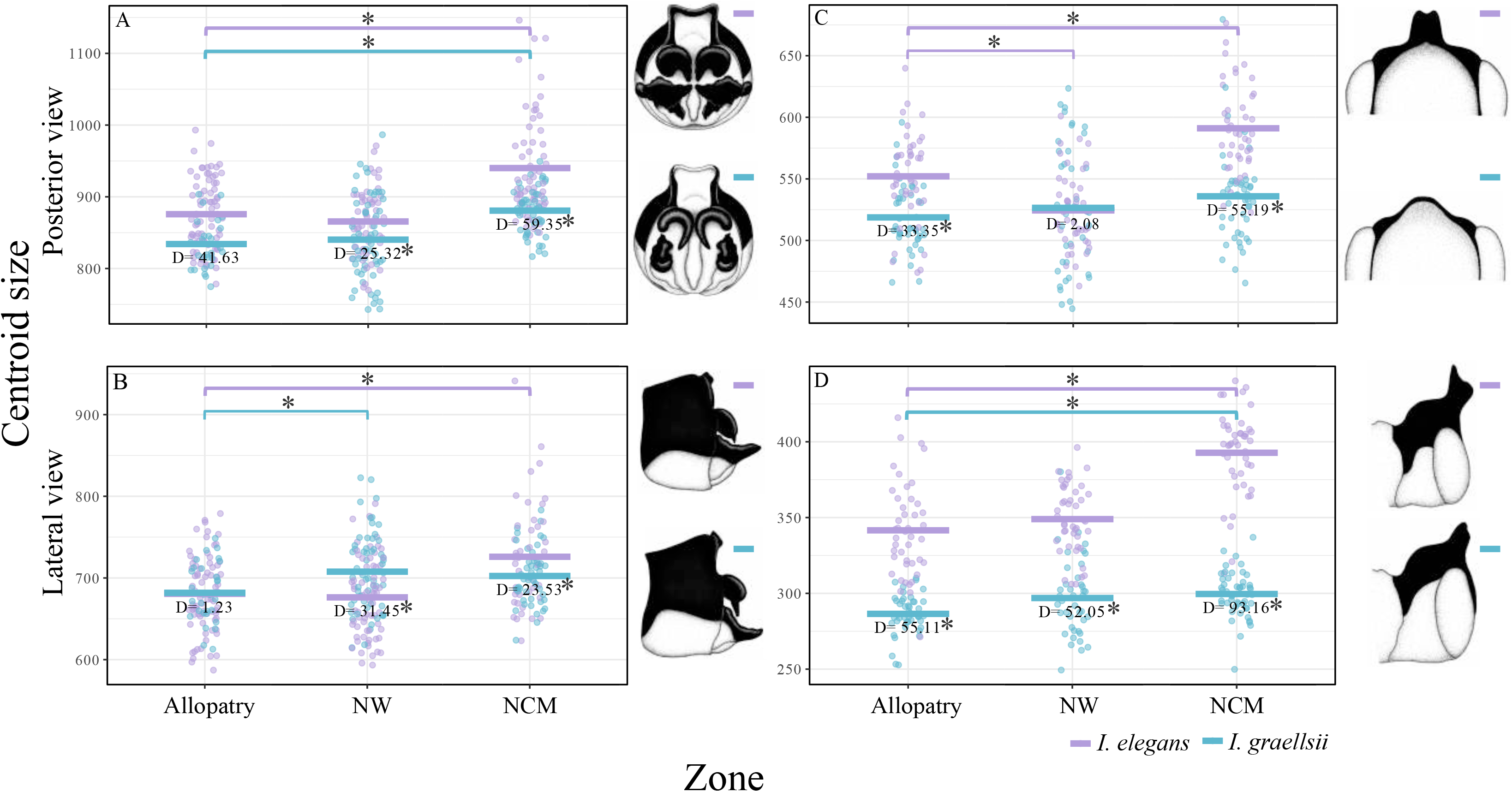
Media values of CS of the male caudal appendages of *I. elegans* and *I. graellsii* in A) posterior and B) lateral views and female prothorax of *I. elegans* and *I. graellsii* in C) posterior and B) lateral views from allopatry and both hybrid zones. Solid lines indicate statistically significant differences at a significant level of p<0.05 between conspecific males between zones.

To investigate whether the detected RCD was unidirectional or bidirectional, i.e., present in one or both species, we conducted comparisons within each species. In *I. elegans*, the CS distance was significantly different among the three zones (posterior view: F=31.345, *p*<0.001; lateral view F=15.194, *p*<0.001; respectively, Table S9; Figure 3A-B). Post hoc analysis showed that the caudal appendages of males from the NCM hybrid zone were significantly larger than those from allopatry (posterior view: z=4.146, p<0.001, d=64.323; lateral view: z=3.444, p<0.001, d=45.350; Table S10), while the males from the NW hybrid zone were not significantly different compared to males from allopatry (Table S10). Similarly, in *I. graellsii*, the CS distance was significantly different among the three zones (posterior view: F=14.533, *p*<0.001; lateral view F=4.080, *p*=0.011; Table S9; Figure 3A-B). Post hoc analysis showed that the caudal appendages of males from the NCM hybrid zone were significantly larger than those from allopatry (posterior view: z=3.265, *p*<0.001, d=46.610; lateral view: z=1.809, *p*=0.023, d=20.588, Table S10), while the males from the NW hybrid zone were only significantly larger than those from allopatry for the lateral view (z=2.219, *p*=0.003, d=25.891). Therefore, RCD (both views) was unilateral, as it was only detected in *I. elegans* from the NCM hybrid zone, while although RCD (lateral view) was also unilateral in the NW hybrid zone, it was only detected in the *I. graellsii* males. (Table 2).

#### Shape variation of female Prothorax

PCA plots of the two views (posterior and lateral) showed a clear separation between species in the three zones (Table S11, Figure 2G-L). Procrustes ANOVA showed statistically significant differences between *I. elegans* and *I. graellsii*: from the allopatric zone (posterior view: F=45.47, *p*<0.001, d=0.12 and lateral view: F=21.21, *p*<0.001, d=0.19); from the NCM hybrid zone (posterior view: F =106.19, p<0.001, d=0.22 and lateral view: F=15.02, p<0.001, d=0.30); and from the NW hybrid zone (posterior view: F=144.23, *p*<0.001, d=0.19 and lateral view: F=42.90, *p*<0.001, d=0.27) (Tables S12-S13). Distance variance analysis confirmed RCD in both views and hybrid regions the NW and the NCM, as distance variance was larger than that in the allopatric zone (distance sympatry - distance allopatry 0) (Table 2).

Within species in *I. elegans,* PC axis1 explained 62.9% (posterior view) and 43.6% (lateral view), and PC axis 2 the 12.4% (posterior view) and 21.7% (lateral view) of the shape variance. They were mainly discriminated in the first axis (Supplementary Figure S4). Procrustes ANOVA showed statistically significant differences between zones (posterior view: F=15.83, *p*<0.001); lateral view: F=13.80, *p*<0.001)] (Table S14). Post hoc paired tests showed statistically significant differences between *I. elegans* individuals from the allopatric zone and the NCM hybrid zone (posterior view: *p*<0.001; d=0.08, and lateral view: *p*<0.001; d=0.11), and between *I. elegans* individuals from the allopatric and the NW hybrid zone at the lateral view (posterior view: *p*=0.486; d=0.03; lateral view: *p*=0.036; d=0.06) (Table S15). Similarly, within species in *I. graellsii*, PC axis1 explained 34.1% (posterior view) and 43.7% (lateral view), and PC axis 2 the 29.3% (posterior view) and 25.1% (lateral view) of the shape variance. They were mainly discriminated in the first axis (Supplementary Figure S4). Procrustes ANOVA showed statistically significant differences between the three zones (Posterior view: F=2.27, *p*=0.008 and lateral view: F=2.18, *p*=0.026) (Table S14). Post hoc paired tests only detected statistically significant differences between individuals of *I. graellsii* from the allopatric and NW hybrid zone in the lateral view (posterior view: *p*=0.470; d=0.02 lateral view: *p*=0.011; d=0.05, Table S15). Therefore, RCD (both views) was unilateral, as it was only detected in *I. elegans* from the NCM hybrid zone, while RCD (lateral view) was bilateral in the NW hybrid zone (Table 2).

#### CS variation of female Prothorax

Procrustes ANOVA showed statistically significant differences between *I. elegans* and *I. graellsii* from allopatry (posterior view: F=24.228, *p*<0.001, d=33.346, and lateral view: F=122.45, *p*<0.001, d=55.109), from the NCM hybrid zone (posterior view: F=54.319, *p*<0.001, d=55.194, and lateral view: F=415.07, *p*<0.001, d=93.158), and from the NW hybrid zone (posterior view: F=0.052, *p*<0.811, d=2.083, lateral view: F=100.34, *p*<0.001, d=52.052) (Tables S16-S17, Figure 3C-D). Distance variance analysis confirmed RCD in both views only in the NCM hybrid zone, where the difference between sympatry and allopatry distances was greater than zero. In contrast, in the NW hybrid zone, the difference between sympatry and allopatry distances was less than zero (Table 2).

Within species, in *I. elegans,* the CS distance of the of female prothoraxes from the NCM hybrid zone was significantly larger than those from allopatry (posterior view: z=3.389, *p*<0.001, d=38.990; lateral view: z=4.357, *p*<0.001, d=51.163; Table S18). Within species, in *I. graellsii,* although the CS distance in the lateral view of females from the NCM hybrid zone (lateral view: z=2.607, *p*<0.001, d=13.115) was significantly larger than those of females from allopatry, they shifted in the same direction than *I. elegans* did, i.e., so that I. graellsii females did not contribute to RCD. In summary, RCD (both views) was unilateral, only detected in *I. elegans* females from the NCM hybrid zone (Table 2).

### Reproductive character displacement at the local scale

Consistently with our fourth prediction that RCD will vary in its strength between populations because of differences in local species proportions (see Sánchez-Guillén et al. 2023). Results are given by shape or CS, sexes, and views (Figure 4).

**Figure 4.**
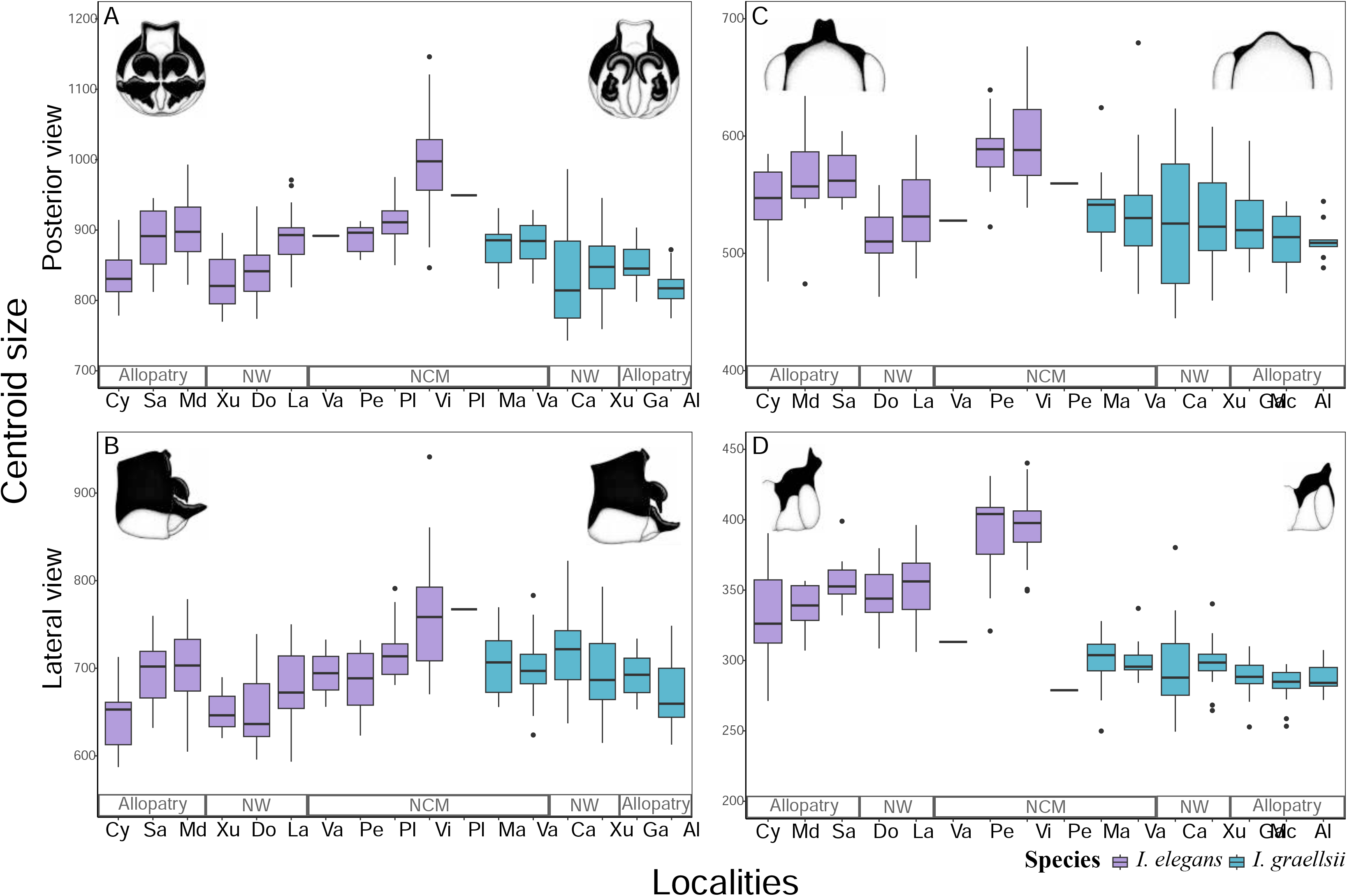
Boxplot representing the morphological variation of CS among studied populations. A) posterior view of male caudal appendage, B) lateral view of male caudal appendage, C) posterior view of female prothorax, D) lateral view of female prothorax. Colors indicate species: purple= *I. elegans*, blue= *I. graellsii*.

#### Shape variation of the male caudal appendages

Procrustes ANOVA applied to assess the shape of the male caudal appendages showed statistically significant differences between localities of both species and in both views: *I. elegans* posterior view: F=5.840, *p*<0.001, and lateral view: F=5.704, *p*<0.001; and *I. graellsii* posterior view: F=4.166, *p*<0.001, and lateral view: F=3.176, *p*<0.001 (Table S19). Posthoc pairwise comparisons between sympatric and allopatric populations of *I. elegans*, showed statistically significant differences for two localities in the NW zone (Doniños and Laxe) and for three localities in the NCM zone (Perdiguero, Valpierre, and Villar) in both the posterior and lateral views, except for Valpierre in the lateral view. In *I. graellsii*, posthoc pairwise comparisons showed statistically significant differences for two localities in the NW zone (Cachadas and Xuño) and for three localities in the NCM zone (Mateo, Plaiaundi, and Valpierre) in both the posterior and lateral views (Table S20).

#### Shape variation of female prothorax

Procrustes ANOVA for the shape of the female prothorax showed statistically significant differences between localities of both species, *I. elegans* (posterior view: F=6.471, *p*<0.001, and lateral view: F=7.248, *p*<0.001) and *I. graellsii* (posterior view: F=1.786, *p*=0.004, and lateral view: F=1.798, *p*=0.016) (Table S21). In *I. elegans* posthoc pairwise comparisons (sympatric populations *versus* allopatric populations) showed statistically significant differences for three out of the four localities from the NCM hybrid zone (posterior view: Perdiguero and Villar; and lateral view: Valpierre, Perdiguero and Villar), and the two localities from the NW hybrid zone (lateral view: Doniños and Laxe). In *I. graellsii*, posthoc pairwise comparisons showed statistically significant differences for two localities from the NW hybrid zone (posterior view: Cachadas and Xuño; lateral view: Cachadas) and two localities from NCM hybrid zone (posterior view: Valpierre and Mateo; lateral view: Mateo) (Table S22).

#### CS variation of male caudal appendages

Procrustes ANOVA for the CS of the male caudal appendages showed statistically significant differences between localities of both species *I. elegans* (posterior view: F=25.352, *p*<0.001, and lateral view: F=14.13, *p*<0.001) (Table S23) and *I. graellsii* (posterior view: F=6.930, *p*<0.001, and lateral view: F=3.083, *p*=0.004) (Table S23). In *I. elegans*, posthoc pairwise comparisons (sympatric populations *versus* allopatric populations) showed statistically significant differences for three out of the four localities from the NCM hybrid zone (both views: Villar, Perdiguero, and Plaiaundi). In *I. graellsii*, posthoc pairwise comparisons showed statistically significant differences for the three populations from the NCM hybrid zone (posterior view: Valpierre, Mateo and Plaiaundi), (Table S24, Figure 4).

#### CS variation of female prothorax

Procrustes ANOVA for the CS of the female prothorax showed statistically significant differences between localities of both species *I. elegans* (posterior view: F=15.355, *p*<0.001, and lateral view: F=16.460, *p*<0.001) (Table S25) and *I. graellsii* (posterior view: F=0.933, *p*=0.469, and lateral view: F=2.213, *p*=0.044) (Table S25). In *I. elegans*, posthoc pairwise comparisons (sympatric populations *versus* allopatric populations) showed statistically significant differences for two localities from the NCM hybrid zone (both views: Villar, Perdiguero), (Table S26, Figure 4). In *I. graellsii*, posthoc pairwise comparisons (sympatric populations *versus* allopatric populations) showed non-statistically significant differences for any population from the NCM hybrid zone.

### Evidence of the lock-and-key coevolution

Consistently with our prediction that reinforcement selection which is stabilizing force, will promote allometric relationships between body size and genital size, we detected a positive allometry between the CS (posterior view) of the caudal appendages of the *I. graellsii* males and the proxy to body size in the NW (Cachadas: r =0.851; *p* =0.001) and in the NCM hybrid zone (Valpierre: r = 0.636; *p* = 0.047), and between the CS (posterior view) of the *I. elegans* females prothorax and the proxy to body size in the NW (Laxe: r=0.657; *p*=0.039), and in the NCM (Perdiguero: r=0.749; *p*=0.012) hybrid zone. However, in the allopatric populations that correlations were not detected (Table S27).

Consistently with our prediction of the lock-and-key coevolution between the male and female genital, we detected moderate coevolution between male caudal appendages and female prothorax in allopatry, but this correlation was slightly lower in both hybrid zones, probably because the different force of the reinforcement and the lock-and key. In detail, for the CS we detected a moderate percentage of coevolution in allopatry: 19% and 20%, posterior and lateral views, respectively in *I. elegans*, and 19% and 4% posterior and lateral views, respectively in *I. graellsii.* However, in both hybrid zones, especially in the NCM, that correlations were considerably reduced: in the NW hybrid zone, 13% and 2%, posterior and lateral views, respectively in *I. elegans*, and 6% and 5% posterior and lateral views, respectively in *I. graellsii;* in the NCM hybrid zone, 7% and 2%, posterior and lateral views, respectively in *I. elegans*, and 1% and 6% posterior and lateral views, respectively in *I. graellsii.* However, for the shape we detected a low percentage of coevolution in allopatry: 3% and 5%, posterior and lateral views, respectively in *I. elegans*, and 7% and 5% posterior and lateral views, respectively in *I. graellsii.* However, in both hybrid zones, especially in the NW, that correlations were slightly increased: in the NW hybrid zone, 6% and 10%, posterior and lateral views, respectively in *I. elegans*, and 6% and 4% posterior and lateral views, respectively in *I. graellsii;* in the NCM hybrid zone, 5% and 5%, posterior and lateral views, respectively in *I. elegans*, and 1% and 6% posterior and lateral views, respectively in *I. graellsii* (Table S28).

## DISCUSSION

The main criterion for RCD is that divergence in reproductive structures is greater in sympatry than in allopatry (see Brown and Wilson 1956).

Divergence in reproductive traits via sexual selection may play a key role in determining the crossability between incipient species (Panhuis et al. 2001; Ritchie 2007). While male genitalia have traditionally been recognized as the most variable and divergent morphological structure, a growing body of empirical evidence supports coevolutionary processes, a pattern of divergent evolution of female’s genitalia matching males’ variation (Simmons and Garcia-Gonzalez 2011, Macagno et al., 2011; Simmons and Fitzpatrick, 2019). Mechanical isolation hypothesis predicts that physical incompatibility between primary genitalia of males and females hinders hybridization (so-called ‘‘lock-and-key’’ or ‘‘structural mechanical isolation’’, Dufour, 1848; Coyne and Orr 2004). Contact and grasping structures that are not intromissive have been included in the category of genital traits (Brennan 2016). In damselflies, species recognition takes place during tandem via tactile interactions, and the female can then accept or refuse to cooperate with the male (Wellenreuther and Sánchez-Guillén, 2016). Moreover, male caudal appendages and female prothorax must match to form a tandem, thus any change not only in size but also in shape will affect the mismatch of these structures. Considering the complexity of these structures we analyzed both the shape and the CS in two complementary views (posterior and lateral) to capture most of the variation.

We directly measured the lock-and-key genital coevolution (shape and CS, and both views) of the complementary male caudal appendages and female prothorax in allopatry as well as both hybrid zones, although coevolution should not be limited to morphological correspondence (Brennan and Prum 2015). We found that while the proportion of interactions showing a positive correlation between the CS of male and female structures was moderate in allopatry (19-20%), it was higher than in each hybrid zone. Note that only the CS analysis (not shape analysis) was useful to investigate the mechanism of lock-and key RI because RCD of the shape was only detected in females. Moreover, consistent with our prediction that genital coevolution results from reinforcing natural selection which represents a stabilizing force that acts to strengthen RI between species, and as a result, it is expected to promote allometric relationships between body size and genital size, we detected a positive allometry between the CS of the *I. graellsii* males and the *I. elegans* females from both hybrid zones.

Our results showed that the more variation was analyzed, the more likely RCD was detected: while the CS was useful to find RCD in both males’ caudal appendages and females’ prothorax, the shape was only useful to find RCD on the female’s prothorax. In a similar manner, the analyses of the two different views of the males’ and females’ genital structures were useful to test the prediction of RCD: while both views (lateral and posterior) were useful to find RCD on the shape of the females’ prothorax, only the lateral view was useful to find RCD on the CS of the males’ caudal appendages and the females’ prothorax. By other hand, the use of geometric morphometrics was opportune since odonate males usually tend to show a pattern of lower allometry than females, i.e., the genital size changes little relative to body size (Nava-Bolaños et al. 2014), which was recovered from our statistical analyses.

Reinforcing selection in response to maladaptive hybridization in hybrid zones can lead to a pattern of RCD of traits involved in premating isolation in sympatry (Howard 1993; Coyne and Orr 2004). However, ecological changes in habitat or resource use may generate changes in the species that also promote RCD (Pfennig and Pfennig, 2009). We detected that both environmental and geographic factors can in part explain several detected patterns of variation in the CS and shape in both hybrid zones and species. Interestingly, the RCD detected of the shape of the *I. graellsii* females was only explained by both geographic factors, latitude, and longitude. Thus, our results highlight the importance of investigating environmental and geographic factors when testing RCD.

RCD pattern can be asymmetric (occurring only in one species or being more pronounced in one of the involved species/sexes) or symmetric (in both involved species). The direction of this pattern depends on the strength of selection to avoid hybridization over each species. As we predicted, based on the reinforcement of the mechanical isolation to form the tandem position in the reciprocal cross direction between *I. graellsii* males and *I. elegans* females (see Arce-Valdés et al., 2023), controlling for environmental and geographic factors we detected that in the NW hybrid zone, CS of the caudal appendages of *I. elegans* and *I. graellsii* males differed more than in allopatry, mainly due to an increase in these structures among the *I. graellsii* males indicating unidirectional RCD, and that shape of the prothorax of *I. elegans* and *I. graellsii* females differed more than in allopatry, resulting from changes in females of both species indicating bidirectional RCD.

Consistently with the prediction that the signal of reinforcement may diminish rapidly once reinforcement ceases to operate, a stronger signal of RCD was detected in the younger NCM hybrid zone than in the older NW hybrid region: CS of the male caudal appendages as well as CS and shape of the female prothorax of *I. elegans* and *I. graellsii* differed more than in allopatry. These differences were due to the increase of these structures in both males and females, which was consistent with the lock-and-key coevolution, but only of *I. elegans* indicating unidirectional RCD. For instance, the beetles *Odontolabis mouhoti* and *O. cuvera* show more divergence in body size, genitalia length and coloration in sympatry than in allopatry; in this system, both species have shifted their relative sizes in opposite directions, i.e., in a bilateral pattern of RCD (Kawano, 2003). However, the size of genital traits of two closely related *Ohomopterus* species differed between populations in contact, however, although both male and female of *C. maiyasanus* increased in sympatry with *C. iwawakianus*, those of *C. iwawakianus* did not differ between allopatric and sympatric populations, indicating a unilateral RCD and providing novel evidence for the lock-and-key hypothesis of genital evolution in beetles (Nishimura et al., 2022).

Asymmetric patterns of RCD arise from different factors such as the asymmetric reinforcement of the reproductive isolation (cf. Cooley, 2007; Yikweon, 2008), the abundance of both parental species, since rarer species have a greater chance of encountering/mating with the common species than *vice versa* (Peterson et al., 2005; Hochkirch et al., 2007), and the level of gene flow with the parental populations (homogenization by gene flow) (Liou and Price 1994; Pfennig and Ryan 2006). When *I. elegans* and *I. graellsii* came into secondary contact, natural selection to avoid hybridization may be stronger for the reciprocal cross between *I. graellsii* males and *I. elegans* females. This is because prezygotic barriers allow hybrid formation, and hybrids experience more disadvantages than the parental species (Arce-Valdés et al, 2023). In contrast, the formation of hybrids from the reciprocal cross between *I. elegans* males and *I. graellsii* females from allopatry is almost completely precluded by gametic isolation (Arce-Valdés et al, 2023). In fact, in the NW hybrid zone (see Arce-Valdés et al, 2023) and the NCM hybrid zone (personal observation) after at least 50 years of sympatry, the strength of the reciprocal crosses between *I. graellsii* males and *I. elegans* females has been enhanced, to the point to be almost completely prevented by the increase of the incompatibility between the male caudal appendages and the female prothorax to form the tandem position. On the contrary, this has not occurred in the reciprocal cross between *I. elegans* males and *I. graellsii* females, among whom the intensity of mechanical isolation is similar between allopatry and sympatry, but the intensity of the postzygotic isolation is lower in sympatry than in allopatry, possibly due to the purge of Bateson–Dobzhansky–Muller (BDM) incompatibilities in sympatry (see Arce-Valdés et al, 2023).

Additionally, *I. elegans* is probably the rarer species in each new Spanish hybrid locality because the secondary contact between *I. elegans* and *I. graellsii* is the result of the range expansion of *I. elegans* over the *I. graellsii* distribution in Spain, so that, the rarer *I. elegans* females have a greater chance of encountering with the common *I. graellsii* males than *vice versa,* and thus be involved in heterospecific matings (Sánchez-Guillén et al. 2023; Arce-Valdés et al, 2023). For instance, *Pterostichus thunbergi* and *P. habui* beetles show unilateral RCD pattern of genital morphology. *Pterostichus thunbergi* differed more from *P. habui* at sympatry than at allopatry, likely because of *P. habui* arrives at the contact zone in small numbers, and thus, would experience stronger selection pressures than *P. habui* (Kosuda et al., 2016). Additionally, interspecific gene flow between *I. elegans* and *I. graellsii* can also explain the increase in the CS of male caudal appendages in *I. graellsii* from the NW hybrid zone, and thus, explain the reduction of the distance variance between *I. elegans* and *I. graellsii* males, because of caudal appendages of hybrids are intermediate between those of the two species (Monetti et al. 2002). In fact, the measured genetic divergence between *I. elegans* and *I. graellsii* in NW hybrid zone was lower than in the NCM hybrid zones and allopatry indicating higher level of interspecific gene flow in the NW than in the NCM hybrid zone (Sánchez-Guillén et al. 2023). A similar finding was detected in the *Carabus, C. maiyasanus* and *C. iwawakianus*, in which hybrid genital traits were intermediate to those of the two species, thus explaining elongation of the genital traits of *C. iwawakianus* (Sasabe et al. 2007).

At the local scale, RCD in the CS and shape of male caudal appendages and female prothorax was detected in all localities (except to in the CS of *I. graellsii* females). Both hybrid zones have introgressed populations of each parental species, i.e., bimodal distribution, and hybrid populations with F_1_, F_2,_ and backcrosses, i.e., unimodal distribution (Sánchez-Guillén et al. 2022). According to this, we would expect a higher relative cost of the hybridization, and thus, RCD will more probably be detected in *I. elegans* females than in *I. graellsii* females (see Kyogoku and Wheatcroft, 2020), but because of the mottled pattern of the hybrid zones, the strength and/or direction of the RCD can lack of consistency between hybrid zones and even between localities (Harrison, 1993). Moreover, the outcome of reproductive interference in sympatric localities is density-dependent (Westman et al., 2002; Hochkirch et al., 2007) thus the species at lower density will experience stronger selective pressure as a result of reproductive interference (Okuzaki, 2013).

## Conclusions

Our study presents novel results in nonterritorial damselflies with few visual displays, but species recognition through mechanic-tactile stimuli, and consequently almost random mating attempts among congeneric species in sympatry. We found evidence of RCD and evidence of the lock-and-key hypothesis of genital evolution in *Ischnura elegans and I. graellsii*. RCD was in part explained by environmental and geographic factors, and in part explained by enhanced reproductive isolation between *I. elegans* females and *I. graellsii* males. Our study adds to previous empirical studies from which has emerged a consensus on the taxonomically widespread role of RCD to lessen reproductive interactions between species (see Goldberg and Lande, 2006). Moreover, our study highlights the importance of discarding environmental and geographic factors when testing RCD and using geometric morphometrics to deal with the complexity of reproductive structures.

## Notes

### Competing Interest Statement

The authors have declared no competing interest.

### Summary of Updates

We changed the name of an author. It was incorrectly written in the previous version. We changed the name of "Sandra Milena Ospina-Garces" to "Sandra M. Ospina-Garces".

