## Supporting Information for "Reproductive character displacement explains strengthening of mechanical barriers in damselflies"

**This PDF file includes:**

Figures S1 to S4

Tables S1 to S28

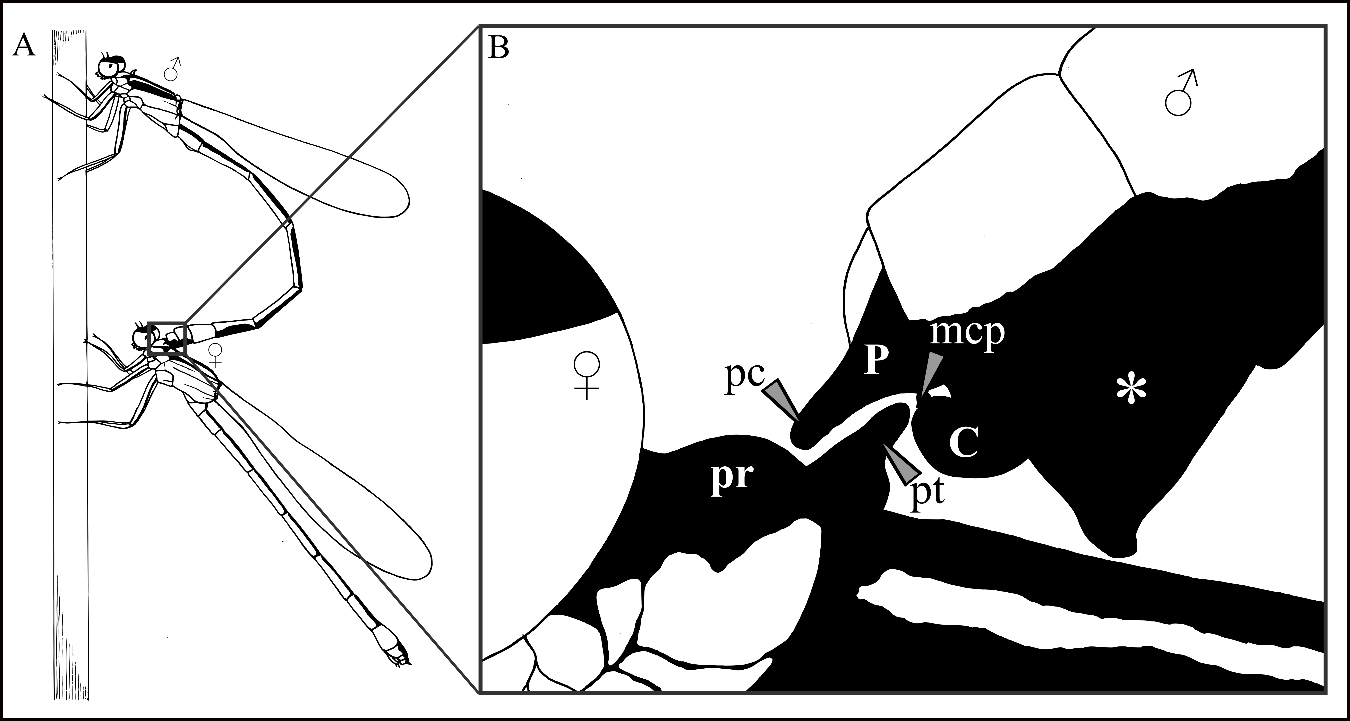

**Figure S1.** A) first contact point in damselflies mating or tandem position: the male grasps the female by her thorax by using his caudal appendages; B) amplification of the contact point: the caudal appendage, located on the last segment of the male´s abdomen, and the pronotum, located on the female´s prothorax. C= cerci, mcp= medio-ventrally directed cercal process, P= paraproct, pc=paraproctal claspers, Pr= prothorax, pt= pronotum *= tenth abdominal segment.

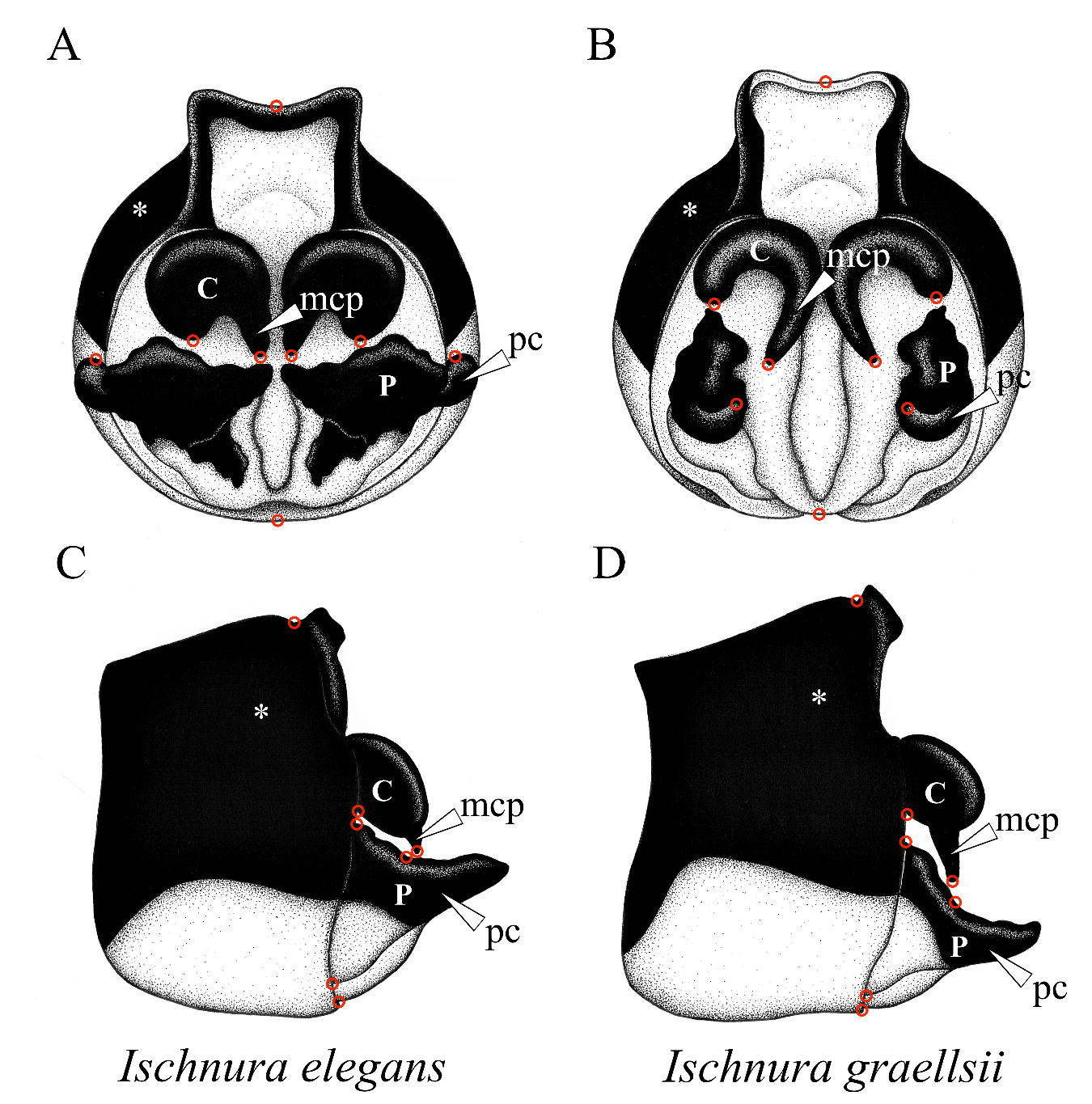

**Figure S2**. Caudal appendage of a male *Ischnura elegans* (A, posterior view and C, left lateral view) and a male of *I. graellsii* (B, posterior view and D, lateral view). C= cerci, mcp= medio-ventrally directed cercal process, P= paraproct, pc= paraproctal claspers, *= tenth abdominal segment. The red circles represent the combination of landmarks that were assigned.

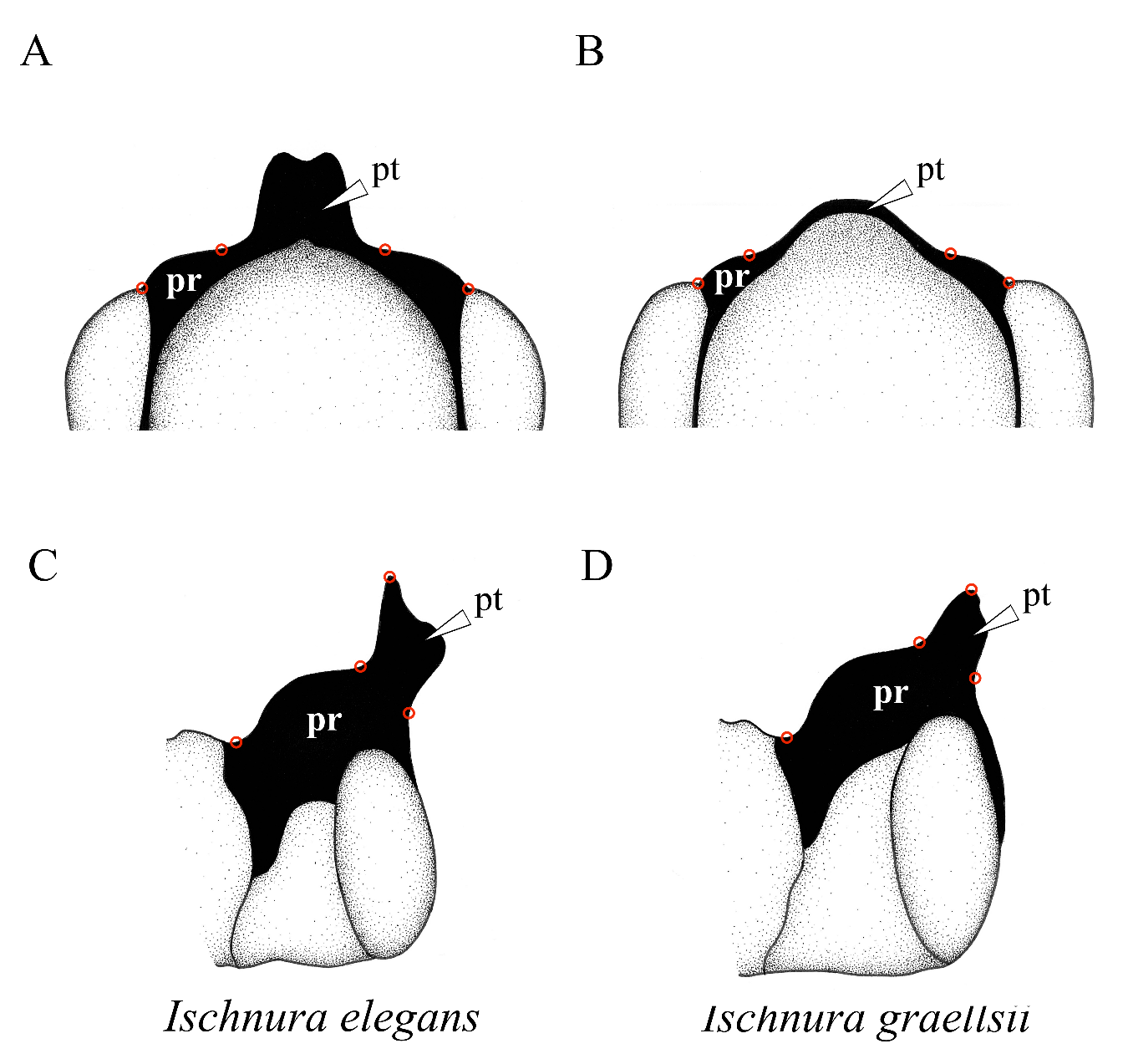

**Figure S3**. Prothorax of a female *Ischnura elegans* (A, posterior view, and C, left lateral view) and a female of *I. graellsii* (B, posterior view, and D, left lateral view). Pr= prothorax, pt= pronotum. The red circles represent the combination of landmarks that were assigned.

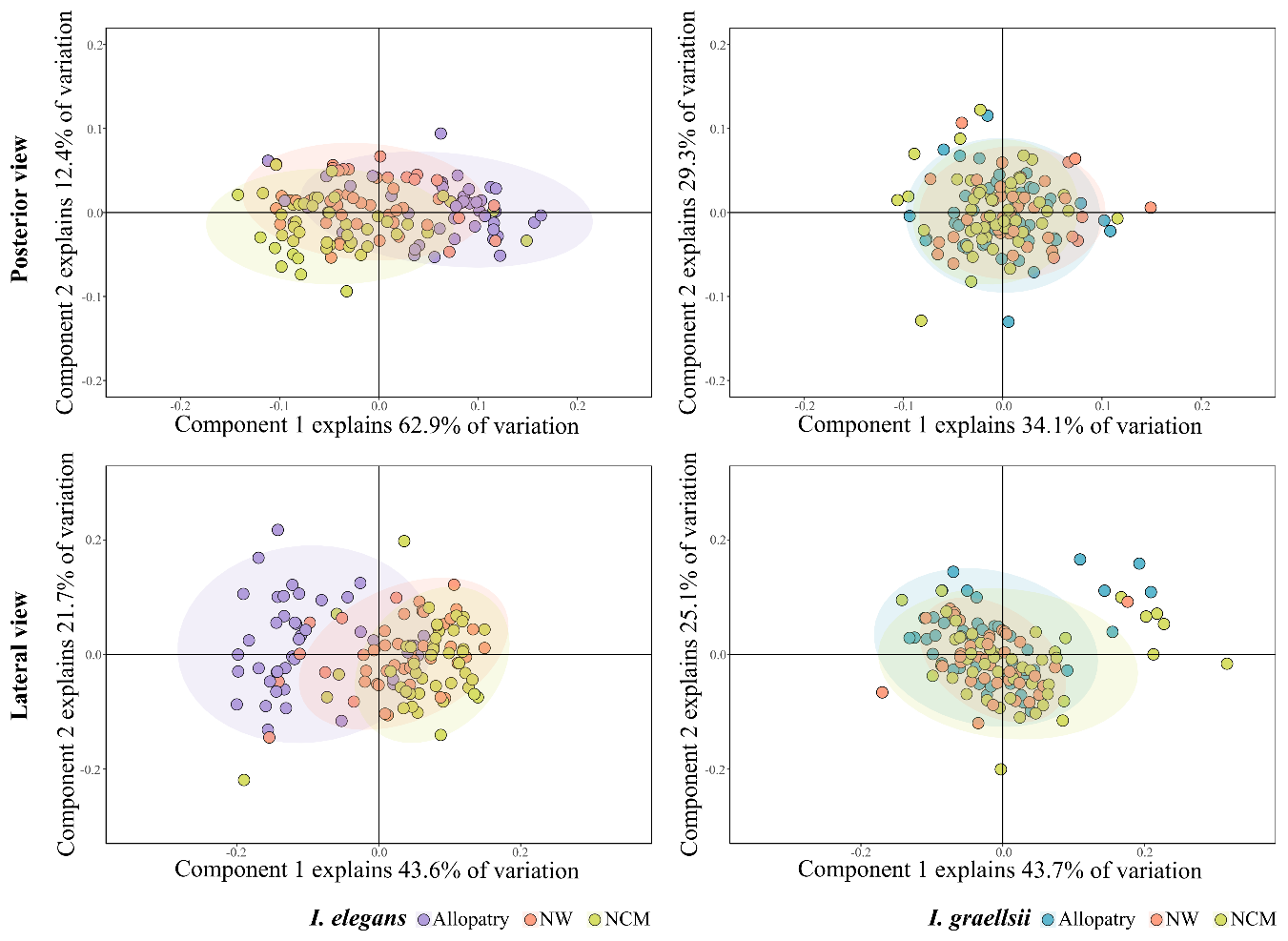

**Figure S4.** Principal Component Analysis (PCA) plots showing the variation of females' caudal appendage of *Ischnura elegans* (A, posterior view: the first two principal axes explain 75.3% of the variance; C, lateral view: the first two principal axes explain 65.3% of the variance) and *I. graellsii* (B, posterior view: the first two principal axes explain 63.4% of the variance; D, lateral view: the first two principal axes explain 68.8% of the variance) in the allopatry region vs. the hybrid regions. NWH= Northwest hybrid region; NCH= North-central hybrid region.

**Table S1**. Landmarks description on reproductive structures involved in the tandem position between *Ischnura elegans* and *I. graellsii*.

| **Character** | **Character view** | **Sex** | **Landmark position** | **Landmark position description** | **Semilandmarks** | **Figure** |
| --- | --- | --- | --- | --- | --- | --- |
| Caudal appendage | Posterior | male | 1 | Tip of the left paraproctal clasper | 80 in four curves: both of cerci. | Figure S2 (A-B) |
|  |  | male | 2 | Tip of the rigth paraproctal clasper |  |  |
|  |  | male | 3 | Tip of the right medio-ventrally directed cercal process |  |  |
|  |  | male | 4 | Tip of the left medio-ventrally directed cercal process |  |  |
|  |  | male | 5 | Most lower point of the left cerci |  |  |
|  |  | male | 6 | Most lower point of the right cerci |  |  |
|  |  | male | 7 | Upper end of tenth abdominal segment |  |  |
|  |  | male | 8 | Lower end of tenth abdominal segment |  |  |
| Caudal appendage | Lateral | male | 1 | Tip of the left medio-ventrally directed cercal process | 36 in two curves: left cerci and left paraproct. | Figure S2 (C-D) |
|  |  | male | 2 | Insertion between the lower part of the left cercus and the tenth abdominal segment |  |  |
|  |  | male | 3 | Insertion between the upper part of the left paraproct and the tenth abdominal segment |  |  |
|  |  | male | 4 | Upper base of left paraproctal clasper |  |  |
|  |  | male | 5 | Insertion between the lower part of the left paraproct and the tenth abdominal segment |  |  |
|  |  | male | 6 | Upper end of the tubercle |  |  |
|  |  | male | 7 | Lower end of the tenth abdominal segment |  |  |
| Prothorax | Posterior | female | 1 | Left most lower point of prothorax | 21 in one curve: pronotum | Figure S3 (A-B) |
|  |  | female | 2 | Left most lower point of pronotum |  |  |
|  |  | female | 3 | Right most lower point of pronotum |  |  |
|  |  | female | 4 | Right most lower point of prothorax |  |  |
| Prothorax | Lateral | female | 1 | Anterior base of prothorax | 19 in one curve: pronotum | Figure S3 (C-D) |
|  |  | female | 2 | Left most lower point of pronotum |  |  |
|  |  | female | 3 | Upper end of the pronotum |  |  |
|  |  | female | 4 | Right most lower point of pronotum |  |  |

| **Table S2.** Results of the correlation test between environmental variables. Variables that presented an absolute value of correlation coefficients (\|r\|) > 0.7 were not included in the same model. | | | | | | |
| --- | --- | --- | --- | --- | --- | --- |
| *I. elegans* males above the diagonal, *I. graellsii* males below the diagonal | | | | |  |  |
| **Latitud** | **-0.11** | 0.79 | **0.41** | **-0.66** | **0.52** | **-0.66** |
| **0.21** | **Longitud** | **-0.31** | **0.007** | 0.76 | **-0.61** | **0.12** |
| **0.55** | **-0.59** | **Precipitation** | **0.46** | -0.78 | **0.66** | -0.71 |
| **0.55** | **-0.61** | 0.97 | **Annual mean temperature** | **-0.39** | 0.77 | -0.94 |
| **-0.62** | **0.62** | -0.93 | -0.94 | **Maximum temperature** | -0.87 | **0.62** |
| **0.49** | -0.72 | 0.95 | 0.98 | -0.98 | **Minimum temperature** | -0.85 |
| **-0.69** | **0.52** | -0.95 | -0.98 | 0.97 | -0.96 | **Elevation** |
| *I. elegans* females above the diagonal, *I. graellsii* females below the diagonal | | | | |  |  |
| **Latitud** | **-0.33** | 0.98 | **0.49** | -0.84 | 0.73 | -0.77 |
| **-0.39** | **Longitud** | **-0.27** | **0.12** | **0.75** | **-0.52** | **0.029** |
| **0.45** | -0.92 | **Precipitation** | **0.4** | -0.77 | **0.62** | -0.7 |
| -0.77 | **0.046** | **0.11** | **Annual mean temperature** | **-0.33** | 0.76 | -0.93 |
| **-0.67** | 0.88 | -0.93 | **0.15** | **Maximum temperature** | -0.84 | **0.58** |
| **-0.27** | **-0.6** | **0.67** | 0.79 | **-0.49** | **Minimum temperature** | -0.85 |
| **0.21** | **0.49** | **-0.64** | -0.78 | **0.47** | -0.97 | **Elevation** |

| **Table S3**. Results of *post-hoc* GLM modeling for the type of distribution (zone) and according to the shape and the CS of male and female reproductive structures. | | | | | | | | | | | | | |
| --- | --- | --- | --- | --- | --- | --- | --- | --- | --- | --- | --- | --- | --- |
| **Morphological character** | **Character view** |  | **Compared zones** |  |  | ***I. elegans*** |  |  | ***I. graellsii*** | | | | |
|  |  |  |  | **Estimate** | **Std. Error** | ***t* value** | ***P*-value** | ***p*<0.05/3** | **Estimate** | **Std. Error** | ***t* value** | ***P*-value** | ***p*<0.05/3** |
| Caudal appendage | Posterior | CS | Allo-NW | -10.157 | 9.991 | -1.017 | 0.311 |  | 6.153 | 10.103 | 0.609 | 0.544 |  |
|  |  |  | Allo-NCM | 64.324 | 9.648 | 6.667 | **<0.001** | * | 46.611 | 10.344 | 4.506 | **<0.001** | * |
|  |  |  | NW-NCM | 74.481 | 10.517 | 7.082 | **<0.001** | * | 40.458 | 8.679 | 4.662 | **<0.001** | * |
|  |  | Shape | Allo-NW | 0.050815 | 0.007298 | 6.963 | **<0.001** | * | 0.038645 | 0.010668 | 3.623 | **<0.002** | * |
|  |  |  | Allo-NCM | 0.035742 | 0.007048 | 5.071 | **<0.001** | * | 0.050661 | 0.010922 | 4.638 | **<0.001** | * |
|  |  |  | NW-NCM | -0.015073 | 0.007683 | -1.962 | 0.0512 |  | 0.012016 | 0.009164 | 1.311 | 0.192 |  |
| Caudal appendage | Lateral | CS | Allo-NW | -16.477 | 8.725 | -1.888 | 0.0605 |  | 24.847 | 9.234 | 2.691 | 0.008 | * |
|  |  |  | Allo-NCM | 43.886 | 8.426 | 5.208 | **<0.001** | * | 21.886 | 9.455 | 2.315 | 0.02214 |  |
|  |  |  | NW-NCM | 60.363 | 9.185 | 6.572 | **<0.001** | * | -2.961 | 7.933 | -0.373 | 0.709 |  |
|  |  | Shape | Allo-NW | 0.071535 | 0.012739 | 5.616 | **<0.001** | * | -0.02251 | 0.012548 | -1.794 | 0.075 |  |
|  |  |  | Allo-NCM | 0.039664 | 0.012301 | 3.224 | **0.001** | * | -0.002359 | 0.012847 | -0.184 | 0.854 |  |
|  |  |  | NW-NCM | -0.031871 | 0.01341 | -2.377 | 0.0185 |  | 0.020151 | 0.010779 | 1.869 | 0.063 |  |
| Prothorax | Posterior | CS | Allo-NW | -27.675 | 7.17 | -3.86 | **<0.001** | * | NA | NA | NA | NA |  |
|  |  |  | Allo-NCM | 38.99 | 7.17 | 5.438 | **<0.001** | * | NA | NA | NA | NA |  |
|  |  |  | NW-NCM | -66.665 | 7.129 | -9.351 | **<0.001** | * | NA | NA | NA | NA |  |
|  |  | Shape | Allo-NW | -0.066664 | 0.013175 | -5.06 | **<0.001** | * | NA | NA | NA | NA |  |
|  |  |  | Allo-NCM | -0.099611 | 0.013175 | -7.56 | **<0.001** | * | NA | NA | NA | NA |  |
|  |  |  | NW-NCM | -0.032947 | 0.013101 | -2.515 | 0.0131 |  | NA | NA | NA | NA |  |
| Prothorax | Lateral | CS | Allo-NW | 7.398 | 5.791 | 1.277 | 0.204 |  | 10.454 | 3.944 | 2.651 | **0.009** | * |
|  |  |  | Allo-NCM | 51.164 | 5.791 | 8.834 | **<0.001** | * | 13.116 | 3.771 | 3.478 | **<0.001** | * |
|  |  |  | NW-NCM | 43.766 | 5.759 | 7.6 | **<0.001** | * | 2.661 | 3.866 | 0.688 | 0.492 |  |
|  |  | Shape | Allo-NW | 0.12364 | 0.0146 | 8.468 | **<0.001** | * | NA | NA | NA | NA |  |
|  |  |  | Allo-NCM | 0.15761 | 0.0146 | 10.794 | **<0.001** | * | NA | NA | NA | NA |  |
|  |  |  | NW-NCM | 0.03397 | 0.01452 | 2.34 | 0.02082 | * | NA | NA | NA | NA |  |

**Table S4**. Variation percentages of the principal component analyzes for the shape of the male caudal appendage between *Ischnura elegans* and *I. graellsii* in each of the regions.

|  | **Allopatry** | | | **Nortwest Hybrid region (NWH)** | | | | **North-central Hybrid region (NCH)** | | |
| --- | --- | --- | --- | --- | --- | --- | --- | --- | --- | --- |
|  | %PC1 | %PC2 | %PC total | %PC1 | %PC2 | %PC total | %PC1 | | %PC2 | %PC total |
| Posterior view | 80.9 | 5.4 | 86.3 | 77.5 | 6.8 | 84.3 | 77.6 | | 5.5 | 83.1 |
| Lateral view | 69.4 | 8.2 | 77.6 | 64 | 11.3 | 75.3 | 61.8 | | 10.8 | 72.6 |

**Table S5**. Results of procrustes ANOVA for shape of male caudal appendages between *Ischnura elegans* and *I. graellsii*

|  | Df | SS | MS | Rsq | F | Z | Pr(>F) |
| --- | --- | --- | --- | --- | --- | --- | --- |
| **Posterior view** |  |  |  |  |  |  |  |
| ***Allopatry*** |  |  |  |  |  |  |  |
| CS | 1 | 0.26269 | 0.26269 | 0.09803 | 46.8192 | 3.7753 | **0.001**** |
| groups | 1 | 1.83282 | 1.83282 | 0.68397 | 326.6667 | 4.8194 | **0.001**** |
| CS:groups | 1 | 0.00627 | 0.00627 | 0.00234 | 1.1181 | 0.4891 | 0.323 |
| Residuals | 103 | 0.5779 | 0.00561 | 0.21566 |  |  |  |
| Total | 106 | 2.67968 |  |  |  |  |  |
| ***North-central hybrid region*** |  |  |  |  |  |  |  |
| CS | 1 | 0.48664 | 0.48664 | 0.18914 | 78.9404 | 4.0982 | **0.001**** |
| groups | 1 | 1.39427 | 1.39427 | 0.54191 | 226.1708 | 4.7433 | **0.001**** |
| CS:groups | 1 | 0.03233 | 0.03233 | 0.01257 | 5.2448 | 3.8148 | **0.001**** |
| Residuals | 107 | 0.65962 | 0.00616 | 0.25638 |  |  |  |
| Total | 110 | 2.57287 |  |  |  |  |  |
| ***Northwest hybrid region*** |  |  |  |  |  |  |  |
| CS | 1 | 0.13857 | 0.13857 | 0.04618 | 20.8936 | 2.9783 | **0.001**** |
| groups | 1 | 2.11415 | 2.11415 | 0.7046 | 318.7827 | 4.4352 | **0.001**** |
| CS:groups | 1 | 0.03818 | 0.03818 | 0.01273 | 5.7574 | 3.5263 | **0.001**** |
| Residuals | 107 | 0.70962 | 0.00663 | 0.2365 |  |  |  |
| Total | 110 | 3.00052 |  |  |  |  |  |
| **Lateral view** |  |  |  |  |  |  |  |
| ***Allopatry*** |  |  |  |  |  |  |  |
| CS | 1 | 0.0691 | 0.0691 | 0.03585 | 9.2434 | 2.5971 | **0.002**** |
| groups | 1 | 1.07839 | 1.07839 | 0.55943 | 144.2526 | 4.6558 | **0.001**** |
| CS:groups | 1 | 0.01017 | 0.01017 | 0.00528 | 1.3606 | 0.8138 | 0.212 |
| Residuals | 103 | 0.77 | 0.00748 | 0.39945 |  |  |  |
| Total | 106 | 1.92766 |  |  |  |  |  |
| ***North-central hybrid region*** |  |  |  |  |  |  |  |
| CS | 1 | 0.06384 | 0.06384 | 0.03754 | 7.2101 | 2.7783 | **0.002**** |
| groups | 1 | 0.79951 | 0.79951 | 0.47009 | 90.2949 | 4.5598 | **0.001**** |
| CS:groups | 1 | 0.03167 | 0.03167 | 0.01862 | 3.5763 | 2.5882 | **0.003**** |
| Residuals | 91 | 0.80576 | 0.00885 | 0.47376 |  |  |  |
| Total | 94 | 1.70078 |  |  |  |  |  |
| ***Northwest hybrid region*** |  |  |  |  |  |  |  |
| CS | 1 | 0.33315 | 0.33315 | 0.14039 | 34.8893 | 3.7696 | **0.001**** |
| groups | 1 | 0.85042 | 0.85042 | 0.35838 | 89.0609 | 4.8738 | **0.001**** |
| CS:groups | 1 | 0.01489 | 0.01489 | 0.00628 | 1.5596 | 1.067 | 0.147 |
| Residuals | 123 | 1.17449 | 0.00955 | 0.49495 |  |  |  |
| Total | 126 | 2.37295 |  |  |  |  |  |

**Table S6**. Results of pairwise comparison of procrustes distances of male caudal appendages estimated between *Ischnura elegans* and *I. graellsii*.

| ***I. elegans* - *I. graellsii* (Posterior view)** | | | | | | | | | | | |
| --- | --- | --- | --- | --- | --- | --- | --- | --- | --- | --- | --- |
| $p.value |  |  |  |  |  | $dist |  |  |  |  |  |
|  | Ie.Allopatry | Ig.Allopatry | Ie.NWH | Ig.NWH | Ie.NCH |  | Ie.Allopatry | Ig.Allopatry | Ie.NWH | Ig.NWH | Ie.NCH |
| Ig.Allopatry | **0.000999** |  |  |  |  | Ig.Allopatry | **0.3136056** |  |  |  |  |
| Ie.NWH | 0.15284715 | **0.000999** |  |  |  | Ie.NWH | 0.06628149 | 0.31704638 |  |  |  |
| Ig.NWH | **0.000999** | 0.41558442 | **0.000999001** |  |  | Ig.NWH | 0.28337955 | 0.04834095 | **0.2870172** |  |  |
| Ie.NCH | 0.18181818 | **0.000999** | 0.393606394 | **0.000999** |  | Ie.NCH | 0.05308722 | 0.30591675 | 0.03954896 | 0.27638172 |  |
| Ig.NCH | **0.000999** | 0.32367632 | **0.000999001** | 0.48751249 | **0.000999** | Ig.NCH | 0.27162187 | 0.06223893 | 0.27418269 | 0.03721039 | **0.2643902** |
| ***I. elegans* - *I. graellsii* (Lateral view)** | | |  |  |  |  |  |  |  |  |  |
|  | Ie.Allopatry | Ig.Allopatry | Ie.NWH | Ig.NWH | Ie.NCH |  | Ie.Allopatry | Ig.Allopatry | Ie.NWH | Ig.NWH | Ie.NCH |
| Ig.Allopatry | **0.000999** |  |  |  |  | Ig.Allopatry | **0.2266591** |  |  |  |  |
| Ie.NWH | 0.16883117 | **0.000999** |  |  |  | Ie.NWH | 0.05738385 | 0.18398247 |  |  |  |
| Ig.NWH | **0.000999** | 0.46053946 | **0.000999001** |  |  | Ig.NWH | 0.24112345 | 0.04609683 | **0.1942156** |  |  |
| Ie.NCH | 0.24875125 | **0.000999** | 0.554445554 | **0.000999** |  | Ie.NCH | 0.05421431 | 0.19031245 | 0.03546307 | 0.20156333 |  |
| Ig.NCH | **0.000999** | 0.71628372 | **0.000999001** | 0.67632368 | **0.000999** | Ig.NCH | 0.22055282 | 0.03419906 | 0.1752149 | 0.0317959 | **0.1818377** |

**Table S7**. Results of procrustes ANOVA for centroid size of male caudal appendages between *Ischnura elegans* and *I. graellsii*.

|  | Df | SS | MS | Rsq | F | Z | Pr(>F) |
| --- | --- | --- | --- | --- | --- | --- | --- |
| **Posterior view** |  |  |  |  |  |  |  |
| ***Allopatry*** |  |  |  |  |  |  |  |
| groups | 1 | 37423 | 37423 | 0.1505 | 18.603 | 3.1243 | **0.001**** |
| Residuals | 105 | 211227 | 2012 | 0.8495 |  |  |  |
| Total | 106 | 248650 |  |  |  |  |  |
| ***North-central hybrid region*** |  |  |  |  |  |  |  |
| groups | 1 | 96780 | 96780 | 0.21346 | 29.581 | 3.9701 | **0.001**** |
| Residuals | 109 | 356618 | 3272 | 0.78654 |  |  |  |
| Total | 110 | 453398 |  |  |  |  |  |
| ***Northwest hybrid region*** |  |  |  |  |  |  |  |
| groups | 1 | 17784 | 17783.6 | 0.05599 | 6.4654 | 1.9986 | **0.013*** |
| Residuals | 109 | 299813 | 2750.6 | 0.94401 |  |  |  |
| Total | 110 | 317597 |  |  |  |  |  |
| **Lateral view** |  |  |  |  |  |  |  |
| ***Allopatry*** |  |  |  |  |  |  |  |
| groups | 1 | 33 | 32.74 | 0.00017 | 0.0177 | -1.3286 | 0.894 |
| Residuals | 105 | 193914 | 1846.8 | 0.99983 |  |  |  |
| Total | 106 | 193947 |  |  |  |  |  |
| ***North-central hybrid region*** |  |  |  |  |  |  |  |
| groups | 1 | 13136 | 13136.1 | 0.05248 | 5.151 | 1.9421 | **0.018*** |
| Residuals | 93 | 237170 | 2550.2 | 0.94752 |  |  |  |
| Total | 94 | 250306 |  |  |  |  |  |
| ***Northwest hybrid region*** |  |  |  |  |  |  |  |
| groups | 1 | 31178 | 31177.5 | 0.09831 | 13.629 | 2.8767 | **0.001**** |
| Residuals | 125 | 285959 | 2287.7 | 0.90169 |  |  |  |
| Total | 126 | 317137 |  |  |  |  |  |

**Table S8**. Results of comparison of centroid size of male caudal appendages between *Ischnura elegans* and *I. graellsii* estimated within regions.

|  | d.obs | UCL (95%) | Zd | Pr(>d) |
| --- | --- | --- | --- | --- |
| **Posterior view** |  |  |  |  |
| ***Allopatry*** |  |  |  |  |
| (Intercept) |  |  |  |  |
| Ig.Allo | 41.63465 | 20.74559 | 2.998461 | **0.001** |
| ***North-central hybrid region*** |  |  |  |  |
| (Intercept) | 880.7223 | 923.9303 | -5.0553 | 1 |
| Ie.NCM | 59.34785 | 23.18013 | 3.72268 | **0.001** |
| ***Northwest hybrid region*** |  |  |  |  |
| (Intercept) |  |  |  |  |
| Ie.NW | 25.32427 | 19.80258 | 1.982883 | **0.013** |
| **Lateral view** |  |  |  |  |
| ***Allopatry*** |  |  |  |  |
| (Intercept) |  |  |  |  |
| Ig.Allo | 1.231523 | 18.75026 | -1.33205 | 0.894 |
| ***North-central hybrid region*** |  |  |  |  |
| (Intercept) | 702.4135 | 721.8293 | -2.25623 | 0.992 |
| Ie.NCM | 23.52983 | 20.20935 | 1.928454 | **0.018** |
| ***Northwest hybrid region*** |  |  |  |  |
| (Intercept) |  |  |  |  |
| Ie.NW | 31.45459 | 17.95901 | 2.807776 | 0.001 |

**Table S9**. Results of procrustes ANOVA for centroid size of male caudal appendages of *Ischnura elegans* and *I. graellsii* between regions.

|  | Df | SS | MS | Rsq | F | Z | Pr(>F) |
| --- | --- | --- | --- | --- | --- | --- | --- |
| ***I. elegans* - Posterior view** |  |  |  |  |  |  |  |
| groups | 2 | 198627 | 99313 | 0.24907 | 31.345 | 5.8315 | **0.001 **** |
| Residuals | 189 | 598834 | 3168 | 0.75093 |  |  |  |
| Total | 191 | 797461 |  |  |  |  |  |
| ***I. elegans* - Lateral view** |  |  |  |  |  |  |  |
| groups | 2 | 79262 | 39631 | 0.13852 | 15.194 | 4.496 | **0.001 **** |
| Residuals | 189 | 492959 | 2608 | 0.86148 |  |  |  |
| Total | 191 | 572221 |  |  |  |  |  |
| ***I. graellsii* - Posterior view** |  |  |  |  |  |  |  |
| groups | 2 | 58310 | 29155 | 0.17825 | 14.533 | 3.9703 | **0.001 **** |
| Residuals | 134 | 268824 | 2006.1 | 0.82175 |  |  |  |
| Total | 136 | 327134 |  |  |  |  |  |
| ***I. graellsii* - Lateral view** |  |  |  |  |  |  |  |
| groups | 2 | 13646 | 6823.1 | 0.0574 | 4.0802 | 2.1344 | **0.011*** |
| Residuals | 134 | 224084 | 1672.3 | 0.9426 |  |  |  |
| Total | 136 | 237730 |  |  |  |  |  |

**Table S10**. Results of pairwise comparison of centroid size of male caudal appendages of *Ischnura elegans* and *I. graellsii* estimated between regions.

|  | d | UCL (95%) | Z | Pr > d |
| --- | --- | --- | --- | --- |
| ***I. elegans* - Posterior view** |  |  |  |  |
| Ie. Allopatry : Ie. NW Hyb | 10.15736 | 23.01598 | 0.316881 | 0.385 |
| Ie. Allopatry : Ie. NCM Hyb | 64.32379 | 21.50365 | 4.146754 | **0.001** |
| Ie. NW Hybrid : Ie. NCM Hyb | 74.48114 | 24.30549 | 4.030365 | **0.001** |
| ***I. elegans* - Lateral view** |  |  |  |  |
| Ie. Allopatry: Ie. NW Hyb | 4.331097 | 16.95851 | -0.37797 | 0.65 |
| Ie. Allopatry: Ie. NCM Hyb | 45.35006 | 19.9127 | 3.444736 | **0.001** |
| Ie. NW Hyb: Ie. NCM Hyb | 49.68116 | 19.96 | 3.49423 | **0.001** |
| ***I. graellsii* - Posterior view** |  |  |  |  |
| Ig. Allopatry: Ig. NW Hyb | 6.153018 | 21.12175 | -0.16506 | 0.565 |
| Ig. Allopatry: Ig. NCM Hyb | 46.61059 | 21.79871 | 3.265549 | **0.001** |
| Ig. NW Hyb: Ig. NCM Hyb | 40.45757 | 18.86424 | 3.149727 | **0.001** |
| ***I. graellsii* - Lateral view** |  |  |  |  |
| Ig. Allopatry: Ig. NW Hyb | 25.89197 | 18.54921 | 2.219969 | **0.003** |
| Ig. Allopatry: Ig. NCM Hyb | 20.58871 | 18.10035 | 1.809825 | **0.023** |
| Ig. NW Hyb: Ig. NCM Hyb | 5.303262 | 15.44343 | 0.018189 | 0.515 |

**Table S11**. Variation percentages of the principal component analyzes for the shape of the female prothorax between *Ischnura elegans* and *I. graellsii* in each of the regions.

|  | **Allopatry** | | | **Nortwest Hybrid region (NWH)** | | | **North-central Hybrid region (NCH)** | | |
| --- | --- | --- | --- | --- | --- | --- | --- | --- | --- |
|  | %PC1 | %PC2 | %PC total | %PC1 | %PC2 | %PC total | %PC1 | %PC2 | %PC total |
| Posterior view | 62.9 | 14.7 | 77.6 | 71.5 | 10.4 | 81.9 | 76.7 | 8.1 | 84.8 |
| Lateral view | 45 | 21.1 | 66.1 | 69 | 11 | 80 | 63.2 | 16.8 | 80 |

**Table S12**. Results of procrustes ANOVA for shape of female prothorax between *Ischnura elegans* and *I. graellsii.*

|  | Df | SS | MS | Rsq | F | Z | Pr(>F) |
| --- | --- | --- | --- | --- | --- | --- | --- |
| **Posterior view** |  |  |  |  |  |  |  |
| ***Allopatry*** |  |  |  |  |  |  |  |
| CS | 1 | 0.18637 | 0.186373 | 0.18294 | 29.6875 | 4.3062 | **0.001**** |
| groups | 1 | 0.28551 | 0.285508 | 0.28025 | 45.4787 | 4.7238 | **0.001**** |
| CS:groups | 1 | 0.01326 | 0.013264 | 0.01302 | 2.1128 | 1.4862 | 0.074 |
| Residuals | 85 | 0.53362 | 0.006278 | 0.52379 |  |  |  |
| Total | 88 | 1.01876 |  |  |  |  |  |
| ***North-central hybrid region*** | |  |  |  |  |  |  |
| CS | 1 | 0.52044 | 0.52044 | 0.31118 | 89.4393 | 4.3856 | **0.001**** |
| groups | 1 | 0.61795 | 0.61795 | 0.36949 | 106.1971 | 5.1823 | **0.001**** |
| CS:groups | 1 | 0.01037 | 0.01037 | 0.0062 | 1.7814 | 1.1681 | 0.124 |
| Residuals | 90 | 0.5237 | 0.00582 | 0.31313 |  |  |  |
| Total | 93 | 1.67246 |  |  |  |  |  |
| ***Northwest hybrid region*** |  |  |  |  |  |  |  |
| CS | 1 | 0.05553 | 0.05553 | 0.04486 | 10.7324 | 2.9993 | **0.001**** |
| groups | 1 | 0.74635 | 0.74635 | 0.60294 | 144.2376 | 4.4285 | **0.001**** |
| CS:groups | 1 | 0.01165 | 0.01165 | 0.00941 | 2.2521 | 1.7484 | **0.044*** |
| Residuals | 82 | 0.4243 | 0.00517 | 0.34278 |  |  |  |
| Total | 85 | 1.23784 |  |  |  |  |  |
| **Lateral view** |  |  |  |  |  |  |  |
| ***Allopatry*** |  |  |  |  |  |  |  |
| CS | 1 | 0.78502 | 0.78502 | 0.31899 | 51.2966 | 6.4805 | **0.001**** |
| groups | 1 | 0.3246 | 0.3246 | 0.1319 | 21.2105 | 5.5705 | **0.001**** |
| CS:groups | 1 | 0.05051 | 0.05051 | 0.02052 | 3.3003 | 2.2343 | **0.007**** |
| Residuals | 85 | 1.30081 | 0.0153 | 0.52858 |  |  |  |
| Total | 88 | 2.46093 |  |  |  |  |  |
| ***North-central hybrid region*** | |  |  |  |  |  |  |
| CS | 1 | 1.8957 | 1.89566 | 0.53948 | 130.4053 | 6.049 | **0.001**** |
| groups | 1 | 0.2183 | 0.21835 | 0.06214 | 15.0205 | 4.8349 | **0.001**** |
| CS:groups | 1 | 0.0915 | 0.09155 | 0.02605 | 6.2977 | 3.1312 | **0.002**** |
| Residuals | 90 | 1.3083 | 0.01454 | 0.37233 |  |  |  |
| Total | 93 | 3.5138 |  |  |  |  |  |
| ***Northwest hybrid region*** |  |  |  |  |  |  |  |
| CS | 1 | 1.17803 | 1.17803 | 0.45385 | 106.673 | 4.5683 | **0.001**** |
| groups | 1 | 0.47384 | 0.47384 | 0.18255 | 42.907 | 5.3718 | **0.001**** |
| CS:groups | 1 | 0.03822 | 0.03822 | 0.01473 | 3.461 | 2.7518 | **0.003**** |
| Residuals | 82 | 0.90556 | 0.01104 | 0.34887 |  |  |  |
| Total | 85 | 2.59565 |  |  |  |  |  |

**Table S13**. Results of pairwise comparison of procrustes distances of female prothorax estimated between *Ischnura elegans* and *I. graellsii.*

| ***I. elegans* - *I. graellsii* (Posterior view)** | | | |  |  |  |  |  |  |  |  |
| --- | --- | --- | --- | --- | --- | --- | --- | --- | --- | --- | --- |
| $p.value |  |  |  |  |  | $dist |  |  |  |  |  |
|  | Ie.Allopatry | Ie.NW | Ie.NCM | Ig.Allopatry | Ig.NW |  | Ie.Allopatry | Ie.NW | Ie.NCM | Ig.Allopatry | Ig.NW |
| Ie.NW | **0.008991** |  |  |  |  | Ie.NWH | 0.09385088 |  |  |  |  |
| Ie.NCM | **0.000999** | 0.2037962 |  |  |  | Ie.NCH | 0.1313733 | 0.04893275 |  |  |  |
| Ig.Allopatry | **0.000999** | **0.000999** | **0.000999001** |  |  | Ig.Allopatry | **0.1256863** | 0.203132 | 0.24002972 |  |  |
| Ig.NW | **0.002997** | **0.000999** | **0.000999001** | 0.95604396 |  | Ig.NWH | 0.11638593 | **0.1910077** | 0.22854328 | 0.01895078 |  |
| Ig.NCM | **0.002997** | **0.000999** | **0.000999001** | 0.82217782 | 0.91808192 | Ig.NCH | 0.10827648 | 0.18849773 | **0.2259387** | 0.02222931 | 0.02125511 |
| ***I. elegans* - *I. graellsii* (Lateral view)** | | |  |  |  |  |  |  |  |  |  |
|  | Ie.Allopatry | Ie.NW | Ie.NCM | Ig.Allopatry | Ig.NW |  | Ie.Allopatry | Ie.NW | Ie.NCM | Ig.Allopatry | Ig.NW |
| Ie.NW | **0.005994** |  |  |  |  | Ie.NWH | 0.13357053 |  |  |  |  |
| Ie.NCM | **0.000999** | 0.46353646 |  |  |  | Ie.NCH | 0.17317678 | 0.05515748 |  |  |  |
| Ig.Allopatry | **0.000999** | **0.000999** | **0.000999001** |  |  | Ig.Allopatry | **0.1976447** | 0.29501254 | 0.32604258 |  |  |
| Ig.NW | **0.000999** | **0.000999** | **0.000999001** | 0.83416583 |  | Ig.NWH | 0.17059242 | **0.272175** | 0.30441633 | 0.03714421 |  |
| Ig.NCM | **0.000999** | **0.000999** | **0.000999001** | 0.51548452 | 0.66633367 | Ig.NCH | 0.17261602 | 0.27230215 | **0.301855** | 0.05135073 | 0.04391936 |

**Table S14**. Results of procrustes ANOVA for the shape of female prothorax of *Ischnura elegans* and *I. graellsii* between regions.

|  |  | Df | SS | MS | Rsq | F | Z | Pr(>F) |
| --- | --- | --- | --- | --- | --- | --- | --- | --- |
|  | ***I. elegans* - Posterior view** | | |  |  |  |  |  |
| CS |  | 1 | 0.09757 | 0.097574 | 0.08266 | 14.6094 | 3.4302 | **0.001 **** |
| groups |  | 2 | 0.21151 | 0.105756 | 0.17918 | 15.8345 | 4.434 | **0.001 **** |
| CS:groups |  | 2 | 0.01649 | 0.008243 | 0.01396 | 1.2341 | 0.6508 | 0.253 |
| Residuals |  | 128 | 0.85489 | 0.006679 | 0.7242 |  |  |  |
| Total |  | 133 | 1.18046 |  |  |  |  |  |
|  | ***I. elegans* - Lateral view** | | |  |  |  |  |  |
| CS |  | 1 | 0.66181 | 0.66181 | 0.22562 | 46.3176 | 6.1663 | **0.001 **** |
| groups |  | 2 | 0.39442 | 0.19721 | 0.13446 | 13.8021 | 6.5019 | **0.001 **** |
| CS:groups |  | 2 | 0.04811 | 0.02405 | 0.0164 | 1.6834 | 1.4398 | 0.068 . |
| Residuals |  | 128 | 1.82893 | 0.01429 | 0.62351 |  |  |  |
| Total |  | 133 | 2.93327 |  |  |  |  |  |
|  | ***I. graellsii* - Posterior view** | | |  |  |  |  |  |
| CS |  | 1 | 0.11016 | 0.110159 | 0.17394 | 28.5362 | 6.3288 | **0.001**** |
| groups |  | 2 | 0.01755 | 0.008774 | 0.02771 | 2.2728 | 2.3783 | **0.008**** |
| CS:groups |  | 2 | 0.00764 | 0.00382 | 0.01206 | 0.9895 | 0.1306 | 0.456 |
| Residuals |  | 129 | 0.49798 | 0.00386 | 0.78629 |  |  |  |
| Total |  | 134 | 0.63333 |  |  |  |  |  |
|  | ***I. graellsii* - Lateral view** | | |  |  |  |  |  |
| CS |  | 1 | 0.26183 | 0.261825 | 0.14506 | 23.4973 | 5.396 | **0.001**** |
| groups |  | 2 | 0.04864 | 0.024319 | 0.02695 | 2.1825 | 1.8806 | **0.026*** |
| CS:groups |  | 2 | 0.05708 | 0.02854 | 0.03162 | 2.5613 | 2.2741 | **0.008**** |
| Residuals |  | 129 | 1.43742 | 0.011143 | 0.79637 |  |  |  |
| Total |  | 134 | 1.80496 |  |  |  |  |  |

**Table S15**. Results of pairwise comparison of procrustes distances for the shape of female prothorax of *Ischnura elegans* and *I. graellsii* estimated between regions.

| ***I. elegans* - Posterior view** | |  |  |  |  |
| --- | --- | --- | --- | --- | --- |
| $p.value |  |  | $dist |  |  |
|  | Ie. Allopatry | Ie. NW hybrid |  | Ie. Allopatry | Ie. NW hybrid |
| Ie. NW hybrid | 0.486 |  | Ie. NW hybrid | 0.02610456 |  |
| Ie. NCM hybrid | **0.001** | **0.004** | Ie. NCM hybrid | 0.08083663 | 0.06240931 |
| ***I. elegans* - Lateral view** | |  |  |  |  |
| $p.value |  |  | $dist |  |  |
|  | Ie. Allopatry | Ie. NW hybrid |  | Ie. Allopatry | Ie. NW hybrid |
| Ie. NW hybrid | **0.036** |  | Ie. NW hybrid | 0.05665246 |  |
| Ie. NCM hybrid | **0.001** | **0.003** | Ie. NCM hybrid | 0.11123438 | 0.06992718 |
| ***I. graellsii* - Posterior view** | |  |  |  |  |
| $p.value |  |  | $dist |  |  |
|  | Ig. Allopatry | Ig. NW Hybrid |  | Ig. Allopatry | Ig. NW Hybrid |
| Ig. NW Hybrid | 0.47052947 |  | Ig. NW Hybrid | 0.01722308 |  |
| Ig. NCM Hybrid | 0.3956044 | **0.04295704** | Ig. NCM Hybrid | 0.01802107 | 0.02340626 |
| ***I. graellsii* - Lateral view** | |  |  |  |  |
| $p.value |  |  | $dist |  |  |
|  | Ig. Allopatry | Ig. NW Hybrid |  | Ig. Allopatry | Ig. NW Hybrid |
| Ig. NW Hybrid | **0.01198801** |  | Ig. NW Hybrid | 0.04752637 |  |
| Ig. NCM Hybrid | 0.14485514 | 0.16583417 | Ig. NCM Hybrid | 0.03242859 | 0.03222597 |

**Table S16**. Results of procrustes ANOVA for centroid size of female prothorax between *Ischnura elegans* and *I. graellsii*.

|  | Df | SS | MS | Rsq | F | Z | Pr(>F) |
| --- | --- | --- | --- | --- | --- | --- | --- |
| **Posterior view** |  |  |  |  |  |  |  |
| ***Allopatry*** |  |  |  |  |  |  |  |
| groups | 1 | 24738 | 24738.4 | 0.21782 | 24.228 | 3.4498 | **0.001**** |
| Residuals | 87 | 88833 | 1021.1 | 0.78218 |  |  |  |
| Total | 88 | 113571 |  |  |  |  |  |
| ***North-central hybrid region*** |  |  |  |  |  |  |  |
| groups | 1 | 71462 | 71462 | 0.37124 | 54.319 | 4.3359 | **0.001**** |
| Residuals | 92 | 121036 | 1316 | 0.62876 |  |  |  |
| Total | 93 | 192499 |  |  |  |  |  |
| ***Northwest hybrid region*** |  |  |  |  |  |  |  |
| groups | 1 | 93 | 93.14 | 0.00062 | 0.0521 | -0.92411 | 0.811 |
| Residuals | 84 | 150255 | 1788.75 | 0.99938 |  |  |  |
| Total | 85 | 150348 |  |  |  |  |  |
| **Lateral view** |  |  |  |  |  |  |  |
| ***Allopatry*** |  |  |  |  |  |  |  |
| groups | 1 | 67566 | 67566 | 0.58462 | 122.45 | 5.6773 | **0.001**** |
| Residuals | 87 | 48006 | 552 | 0.41538 |  |  |  |
| Total | 88 | 115573 |  |  |  |  |  |
| ***North-central hybrid region*** |  |  |  |  |  |  |  |
| groups | 1 | 203574 | 203574 | 0.81857 | 415.07 | 8.2078 | **0.001**** |
| Residuals | 92 | 45122 | 490 | 0.18143 |  |  |  |
| Total | 93 | 248695 |  |  |  |  |  |
| ***Northwest hybrid region*** |  |  |  |  |  |  |  |
| groups | 1 | 58128 | 58128 | 0.54432 | 100.34 | 5.2049 | **0.001**** |
| Residuals | 84 | 48663 | 579 | 0.45568 |  |  |  |
| Total | 85 | 106791 |  |  |  |  |  |

**Table S17**. Results of comparison of centroid size of female prothorax between *Ischnura elegans* and *I. graellsii* estimated within regions.

|  | d.obs | UCL (95%) | Zd | Pr(>d) |
| --- | --- | --- | --- | --- |
| **Posterior view** |  |  |  |  |
| ***Allopatry*** |  |  |  |  |
| (Intercept) |  |  |  |  |
| Ig.Alo | 33.34636 | 15.4369 | 3.234414 | **0.001** |
| ***North-central hybrid region*** |  |  |  |  |
| (Intercept) | 535.8747 | 569.746 | -5.15583 | 1 |
| Ie.NCM | 55.19483 | 19.53223 | 3.829502 | **0.001** |
| ***Northwest hybrid region*** |  |  |  |  |
| (Intercept) |  |  |  |  |
| Ie.NW | 2.083637 | 18.55167 | -0.92877 | 0.811 |
| **Lateral view** |  |  |  |  |
| ***Allopatry*** |  |  |  |  |
| (Intercept) |  |  |  |  |
| Ig.Alo | 55.10968 | 15.39399 | 4.499152 | **0.001** |
| ***North-central hybrid region*** |  |  |  |  |
| (Intercept) | 299.5823 | 352.2892 | -7.11816 | 1 |
| Ie.NCM | 93.1581 | 20.92364 | 5.260871 | **0.001** |
| ***Northwest hybrid region*** |  |  |  |  |
| (Intercept) |  |  |  |  |
| Ie.NW | 52.05294 | 14.68693 | 4.25363 | **0.001** |

**Table S18**. Results of pairwise comparison of centroid size of female prothorax of *Ischnura elegans* and *I. graellsii* estimated between regions. Ie: *Ischnura elegans*, Ig: *I. graellsii.*

|  | d | UCL (95%) | Z | Pr > d |
| --- | --- | --- | --- | --- |
| ***I. elegans* - Posterior view** |  |  |  |  |
| Ie. Allopatry: Ie. NW | 27.67486 | 18.10051 | 2.404988 | **0.004** |
| Ie. Allopatry: Ie. NCM | 38.99031 | 17.64992 | 3.389354 | **0.001** |
| Ie. NW: Ie. NCM | 66.66516 | 17.85096 | 4.845951 | **0.001** |
| ***I. elegans* - Lateral view** |  |  |  |  |
| Ie. Allopatry: Ie. NW | 7.397704 | 14.67006 | 0.520667 | 0.326 |
| Ie. Allopatry: Ie. NCM | 51.16398 | 14.83876 | 4.357244 | **0.001** |
| Ie. NW: Ie. NCM | 43.76628 | 14.57763 | 4.389795 | **0.001** |
| ***I. graellsii* - Posterior view** |  |  |  |  |
| Ig. Allopatry: Ig. NW | 3.212651 | 17.61076 | -0.60264 | 0.72 |
| Ig. Allopatry: Ig. NCM | 15.31244 | 17.80671 | 1.336112 | 0.084 |
| Ig. NW: Ig. NCM | 12.09979 | 16.30323 | 1.085092 | 0.15 |
| ***I. graellsii* - Lateral view** |  |  |  |  |
| Ig. Allopatry: Ig. NW | 10.45445 | 7.869537 | 2.14077 | **0.008** |
| Ig. Allopatry: Ig. NCM | 13.11556 | 7.609655 | 2.607355 | **0.001** |
| Ig. NW: Ig. NCM | 2.661109 | 7.991176 | -0.04195 | 0.528 |

**Table S19**. Results of Procrustes ANOVA for the shape of the male caudal appendages of *Ischnura elegans* and *I. graellsii* between localities.

|  | Df | SS | MS | Rsq | F | Z | Pr(>F) |
| --- | --- | --- | --- | --- | --- | --- | --- |
| ***I. elegans* - Posterior view** |  |  |  |  |  |  |  |
| CS | 1 | 0.06442 | 0.064419 | 0.05013 | 12.6652 | 4.8199 | 0.001 |
| groups | 9 | 0.26736 | 0.029707 | 0.20806 | 5.8406 | 8.1096 | **0.001** |
| CS:groups | 9 | 0.07838 | 0.008709 | 0.061 | 1.7123 | 2.9136 | 0.004 |
| Residuals | 172 | 0.87485 | 0.005086 | 0.68081 |  |  |  |
| Total | 191 | 1.28501 |  |  |  |  |  |
| ***I. elegans* - Lateral view** |  |  |  |  |  |  |  |
| CS | 1 | 0.16541 | 0.165414 | 0.06837 | 17.4099 | 4.3024 | 0.001 |
| groups | 9 | 0.48775 | 0.054195 | 0.2016 | 5.704 | 6.7867 | **0.001** |
| CS:groups | 9 | 0.13199 | 0.014665 | 0.05455 | 1.5435 | 1.8545 | 0.028 |
| Residuals | 172 | 1.6342 | 0.009501 | 0.67547 |  |  |  |
| Total | 191 | 2.41935 |  |  |  |  |  |
| ***I. graellsii* - Posterior view** | |  |  |  |  |  |  |
| CS | 1 | 0.03541 | 0.035413 | 0.03771 | 6.0968 | 3.4264 | 0.001 |
| groups | 6 | 0.14521 | 0.024202 | 0.15464 | 4.1667 | 5.1645 | **0.001** |
| CS:groups | 5 | 0.03813 | 0.007627 | 0.04061 | 1.313 | 1.1069 | 0.138 |
| Residuals | 124 | 0.72025 | 0.005808 | 0.76703 |  |  |  |
| Total | 136 | 0.93901 |  |  |  |  |  |
| ***I. graellsii* - Lateral view** |  |  |  |  |  |  |  |
| CS | 1 | 0.12438 | 0.124377 | 0.12017 | 21.0913 | 4.3332 | 0.001 |
| groups | 6 | 0.11239 | 0.018732 | 0.1086 | 3.1765 | 4.9523 | **0.001** |
| CS:groups | 5 | 0.06696 | 0.013393 | 0.0647 | 2.2711 | 2.9263 | 0.001 |
| Residuals | 124 | 0.73124 | 0.005897 | 0.70653 |  |  |  |
| Total | 136 | 1.03497 |  |  |  |  |  |

**Table S20**. Results of pairwise comparison of procrustes distances for the shape of the male caudal appendages of *Ischnura elegans* and *I. graellsii* estimated between localities. Ie: *Ischnura elegans*, Ig: *I. graellsii*, Al: Alba, Ca: Cachadas, Cy: St. Cyprien, Do: Doniños, Ga: Gamillazo, He: Hervias, La: Laxe, Ma: Mateo, Md: Marais d’Orx, Pe: Perdiguero, Pl: Plaiaundi, Sa: Sabiere, Vi: Villar, Xu: Xuño.

| ***I. elegans* (Posterior view)** | | |  |  |  |  |  |  |  |  |  |  |  |  |  |  |  |  |  |
| --- | --- | --- | --- | --- | --- | --- | --- | --- | --- | --- | --- | --- | --- | --- | --- | --- | --- | --- | --- |
| $p.value |  |  |  |  |  |  |  |  |  | $dist |  |  |  |  |  |  |  |  |  |
|  | Ie.Cy | Ie.Do | Ie.He | Ie.La | Ie.Md | Ie.Pe | Ie.Pl | Ie.Sa | Ie.Vi |  | Ie.Cy | Ie.Do | Ie.He | Ie.La | Ie.Md | Ie.Pe | Ie.Pl | Ie.Sa | Ie.Vi |
| Ie.Do | **0.000999** |  |  |  |  |  |  |  |  | Ie.Do | 0.06610639 |  |  |  |  |  |  |  |  |
| Ie.He | 0.06693307 | 0.76923077 |  |  |  |  |  |  |  | Ie.He | 0.08206711 | 0.0472009 |  |  |  |  |  |  |  |
| Ie.La | **0.000999** | 0.17082917 | 0.515484515 |  |  |  |  |  |  | Ie.La | 0.06157226 | 0.03233694 | 0.0557717 |  |  |  |  |  |  |
| Ie.Md | 0.2947053 | **0.000999** | **0.03996004** | **0.000999** |  |  |  |  |  | Ie.Md | 0.03063287 | 0.07729442 | 0.09020455 | 0.06621183 |  |  |  |  |  |
| Ie.Pe | **0.000999** | **0.001998** | 0.675324675 | **0.024975** | **0.000999** |  |  |  |  | Ie.Pe | 0.06455585 | 0.0551442 | 0.05171117 | 0.04520488 | 0.06710504 |  |  |  |  |
| Ie.Pl | 0.26073926 | **0.000999** | 0.074925075 | **0.000999** | 0.49250749 | **0.000999** |  |  |  | Ie.Pl | 0.03569734 | 0.06575622 | 0.08388618 | 0.06013931 | 0.03168331 | 0.06976739 |  |  |  |
| Ie.Sa | 0.07392607 | **0.000999** | 0.108891109 | **0.000999** | 0.81518482 | **0.000999** | 0.72527473 |  |  | Ie.Sa | 0.03610904 | 0.06723836 | 0.07762808 | 0.05427661 | 0.02297947 | 0.05741847 | 0.02802662 |  |  |
| Ie.Vi | **0.000999** | **0.000999** | 0.458541459 | **0.002997** | **0.000999** | 0.32767233 | **0.000999** | **0.000999** |  | Ie.Vi | 0.06968765 | 0.05319571 | 0.05813361 | 0.04818198 | 0.07406544 | 0.033081 | 0.07119689 | 0.06551219 |  |
| Ie.Xu | 0.18881119 | 0.05294705 | 0.185814186 | 0.07492508 | 0.16683317 | 0.07292707 | 0.52347652 | 0.23276723 | 0.16883117 | Ie.Xu | 0.06593151 | 0.08096019 | 0.09176536 | 0.07713209 | 0.06712778 | 0.07872981 | 0.05662301 | 0.06392137 | 0.06686554 |
| ***I. elegans* (Lateral view)** | | |  |  |  |  |  |  |  |  |  |  |  |  |  |  |  |  |  |
| $p.value |  |  |  |  |  |  |  |  |  | $dist |  |  |  |  |  |  |  |  |  |
|  | Ie.Cy | Ie.Do | Ie.He | Ie.La | Ie.Md | Ie.Pe | Ie.Pl | Ie.Sa | Ie.Vi |  | Ie.Cy | Ie.Do | Ie.He | Ie.La | Ie.Md | Ie.Pe | Ie.Pl | Ie.Sa | Ie.Vi |
| Ie.Do | **0.000999** |  |  |  |  |  |  |  |  | Ie.Do | 0.08658854 |  |  |  |  |  |  |  |  |
| Ie.He | 0.12887113 | 0.22377622 |  |  |  |  |  |  |  | Ie.He | 0.10644455 | 0.09144708 |  |  |  |  |  |  |  |
| Ie.La | **0.01398601** | **0.01198801** | 0.277722278 |  |  |  |  |  |  | Ie.La | 0.05573113 | 0.05674802 | 0.08612442 |  |  |  |  |  |  |
| Ie.Md | **0.02597403** | **0.000999** | 0.102897103 | **0.000999** |  |  |  |  |  | Ie.Md | 0.05479268 | 0.12342413 | 0.11333014 | 0.08371553 |  |  |  |  |  |
| Ie.Pe | 0.11188811 | **0.000999** | 0.435564436 | 0.07992008 | **0.003996** |  |  |  |  | Ie.Pe | 0.04980431 | 0.07758064 | 0.07721929 | 0.0527597 | 0.07417069 |  |  |  |  |
| Ie.Pl | 0.3966034 | **0.002997** | 0.447552448 | 0.08491509 | 0.07792208 | 0.36763237 |  |  |  | Ie.Pl | 0.04126143 | 0.07802408 | 0.07749259 | 0.05588857 | 0.0570798 | 0.04489406 |  |  |  |
| Ie.Sa | 0.23676324 | **0.000999** | 0.115884116 | **0.00499501** | **0.04495505** | **0.01398601** | 0.43956044 |  |  | Ie.Sa | 0.03869053 | 0.09115838 | 0.10725409 | 0.06114974 | 0.05093584 | 0.06162067 | 0.0401753 |  |  |
| Ie.Vi | 0.08691309 | **0.01698302** | 0.201798202 | 0.17682318 | **0.000999** | 0.13286713 | 0.08191808 | **0.001998** |  | Ie.Vi | 0.04755629 | 0.05843811 | 0.09372728 | 0.04102586 | 0.0856495 | 0.04781496 | 0.05601534 | 0.06423278 |  |
| Ie.Xu | 0.14285714 | 0.46653347 | 0.806193806 | 0.24475525 | **0.02797203** | 0.56143856 | 0.37062937 | 0.06793207 | 0.32367632 | Ie.Xu | 0.09457048 | 0.07129573 | 0.07399751 | 0.0831322 | 0.1245708 | 0.06704399 | 0.07855841 | 0.10623719 | 0.07770814 |
| ***I. graellsii* (Posterior view)** | | |  |  |  |  |  |  |  |  |  |  |  |  |  |  |  |  |  |
| $p.value |  |  |  |  |  |  |  |  |  | $dist |  |  |  |  |  |  |  |  |  |
|  | Ig.Al | Ig.Ca | Ig.Ga | Ig.He | Ig.Ma | Ig.Pl |  |  |  |  | Ig.Al | Ig.Ca | Ig.Ga | Ig.He | Ig.Ma | Ig.Pl |  |  |  |
| Ig.Ca | **0.005994** |  |  |  |  |  |  |  |  | Ig.Ca | 0.05155853 |  |  |  |  |  |  |  |  |
| Ig.Ga | 0.73326673 | **0.000999** |  |  |  |  |  |  |  | Ig.Ga | 0.0286857 | 0.06634806 |  |  |  |  |  |  |  |
| Ig.He | **0.000999** | **0.000999** | **0.000999001** |  |  |  |  |  |  | Ig.He | 0.06294259 | 0.05961725 | 0.07363648 |  |  |  |  |  |  |
| Ig.Ma | 0.05594406 | **0.008991** | **0.001998002** | **0.001998** |  |  |  |  |  | Ig.Ma | 0.04189589 | 0.0416085 | 0.05575986 | 0.04438594 |  |  |  |  |  |
| Ig.Pl | **0.046953** | 0.07592408 | **0.01998002** | 0.07492508 | 0.06093906 |  |  |  |  | Ig.Pl | 0.15182619 | 0.12912013 | 0.17134207 | 0.13066339 | 0.13779419 |  |  |  |  |
| Ig.Xu | 0.22677323 | **0.003996** | **0.024975025** | **0.000999** | 0.6033966 | 0.05294705 |  |  |  | Ig.Xu | 0.03278553 | 0.04422558 | 0.04485898 | 0.0563608 | 0.02557397 | 0.14375651 |  |  |  |
| ***I. graellsii* (Lateral view)** | | |  |  |  |  |  |  |  |  |  |  |  |  |  |  |  |  |  |
| $p.value |  |  |  |  |  |  |  |  |  | $dist |  |  |  |  |  |  |  |  |  |
|  | Ig.Al | Ig.Ca | Ig.Ga | Ig.He | Ig.Ma | Ig.Pl |  |  |  |  | Ig.Al | Ig.Ca | Ig.Ga | Ig.He | Ig.Ma | Ig.Pl |  |  |  |
| Ig.Ca | **0.005994** |  |  |  |  |  |  |  |  | Ig.Ca | 0.05155853 |  |  |  |  |  |  |  |  |
| Ig.Ga | 0.73326673 | **0.000999** |  |  |  |  |  |  |  | Ig.Ga | 0.0286857 | 0.06634806 |  |  |  |  |  |  |  |
| Ig.He | **0.000999** | **0.000999** | **0.000999001** |  |  |  |  |  |  | Ig.He | 0.06294259 | 0.05961725 | 0.07363648 |  |  |  |  |  |  |
| Ig.Ma | 0.05594406 | **0.008991** | **0.001998002** | **0.001998** |  |  |  |  |  | Ig.Ma | 0.04189589 | 0.0416085 | 0.05575986 | 0.04438594 |  |  |  |  |  |
| Ig.Pl | **0.046953** | 0.07592408 | **0.01998002** | 0.07492508 | 0.06093906 |  |  |  |  | Ig.Pl | 0.15182619 | 0.12912013 | 0.17134207 | 0.13066339 | 0.13779419 |  |  |  |  |
| Ig.Xu | 0.22677323 | **0.003996** | **0.024975025** | **0.000999** | 0.6033966 | 0.05294705 |  |  |  | Ig.Xu | 0.03278553 | 0.04422558 | 0.04485898 | 0.0563608 | 0.02557397 | 0.14375651 |  |  |  |

**Table S21**. Results of Procrustes ANOVA for the shape of female prothorax of *Ischnura elegans* and *I. graellsii* between localities.

|  | Df | SS | MS | Rsq | F | Z | Pr(>F) |
| --- | --- | --- | --- | --- | --- | --- | --- |
| ***I. elegans* - Posterior view** |  |  |  |  |  |  |  |
| CS | 1 | 0.09757 | 0.097574 | 0.08266 | 15.5696 | 3.5026 | 0.001 |
| groups | 7 | 0.2839 | 0.040557 | 0.2405 | 6.4716 | 5.6177 | **0.001** |
| CS:groups | 6 | 0.05322 | 0.00887 | 0.04508 | 1.4154 | 1.1625 | 0.12 |
| Residuals | 119 | 0.74577 | 0.006267 | 0.63176 |  |  |  |
| Total | 133 | 1.18046 |  |  |  |  |  |
| ***I. elegans* - Lateral view** |  |  |  |  |  |  |  |
| CS | 1 | 0.66181 | 0.66181 | 0.22562 | 52.4416 | 6.3299 | 0.001 |
| groups | 7 | 0.64036 | 0.09148 | 0.21831 | 7.2488 | 8.7361 | **0.001** |
| CS:groups | 6 | 0.12933 | 0.02156 | 0.04409 | 1.708 | 2.1542 | 0.018 |
| Residuals | 119 | 1.50177 | 0.01262 | 0.51198 |  |  |  |
| Total | 133 | 2.93327 |  |  |  |  |  |
| ***I. graellsii* - Posterior view** | |  |  |  |  |  |  |
| CS | 1 | 0.11016 | 0.110159 | 0.17394 | 29.5266 | 6.3595 | 0.001 |
| groups | 7 | 0.04666 | 0.006666 | 0.07368 | 1.7867 | 2.6976 | **0.004** |
| CS:groups | 6 | 0.02881 | 0.004801 | 0.04549 | 1.2869 | 1.1092 | 0.14 |
| Residuals | 120 | 0.4477 | 0.003731 | 0.7069 |  |  |  |
| Total | 134 | 0.63333 |  |  |  |  |  |
| ***I. graellsii* - Lateral view** |  |  |  |  |  |  |  |
| CS | 1 | 0.26183 | 0.261825 | 0.14506 | 24.0024 | 5.4325 | 0.001 |
| groups | 7 | 0.13733 | 0.019619 | 0.07609 | 1.7986 | 2.1436 | **0.016** |
| CS:groups | 6 | 0.09681 | 0.016135 | 0.05363 | 1.4791 | 1.3676 | 0.091 |
| Residuals | 120 | 1.30899 | 0.010908 | 0.72522 |  |  |  |
| Total | 134 | 1.80496 |  |  |  |  |  |

**Table S22**. Results of pairwise comparison of procrustes distances for the shape of female prothorax of *Ischnura elegans* and *I. graellsii* estimated between localities. Ie: *Ischnura elegans*, Ig: *I. graellsii*, Al: Alba, Ca: Cachadas, Cy: St. Cyprien, Do: Doniños, Ga: Gamillazo, He: Hervias, La: Laxe, Ma: Mateo, Mc: Saïdia, Pe: Perdiguero, Sa: Sabiere, Vi: Villar, Xu: Xuño.

| ***I. elegans* (Posterior view)** | | |  |  |  |  |  |  |  |  |  |  |  |  |  |
| --- | --- | --- | --- | --- | --- | --- | --- | --- | --- | --- | --- | --- | --- | --- | --- |
| $p.value |  |  |  |  |  |  |  | $dist |  |  |  |  |  |  |  |
|  | Ie.Do | Ie.Cy | Ie.He | Ie.La | Ie.Md | Ie.Pe | Ie.Sa |  | Ie.Do | Ie.Cy | Ie.He | Ie.La | Ie.Md | Ie.Pe | Ie.Sa |
| Ie.Cy | 0.39160839 |  |  |  |  |  |  | Ie.Cy | 0.03757583 |  |  |  |  |  |  |
| Ie.He | 0.11688312 | 0.35964036 |  |  |  |  |  | Ie.He | 0.13354619 | 0.10313621 |  |  |  |  |  |
| Ie.La | 0.24475525 | 0.88811189 | 0.388611389 |  |  |  |  | Ie.La | 0.04640793 | 0.02373599 | 0.09992092 |  |  |  |  |
| Ie.Md | 0.99200799 | 0.84515485 | 0.1998002 | 0.52647353 |  |  |  | Ie.Md | 0.02167649 | 0.02990451 | 0.12719073 | 0.04110949 |  |  |  |
| Ie.Pe | 0.63836164 | 0.07692308 | **0.041958042** | 0.05294705 | 0.56343656 |  |  | Ie.Pe | 0.03207041 | 0.05984936 | 0.15541049 | 0.07027335 | 0.04062927 |  |  |
| Ie.Sa | 0.05894106 | 0.28471529 | 0.681318681 | 0.4035964 | 0.2017982 | **0.010989** |  | Ie.Sa | 0.08400956 | 0.05499412 | 0.0788099 | 0.04964461 | 0.07489598 | 0.110062 |  |
| Ie.Vi | **0.036963** | **0.003996** | **0.008991009** | **0.000999** | 0.07992008 | 0.27172827 | **0.000999** | Ie.Vi | 0.06783063 | 0.08895477 | 0.17986653 | 0.10019511 | 0.07360233 | 0.0437461 | 0.13887707 |
| ***I. elegans* (Lateral view)** | | |  |  |  |  |  |  |  |  |  |  |  |  |  |
| $p.value |  |  |  |  |  |  |  | $dist |  |  |  |  |  |  |  |
|  | Ie.Do | Ie.Cy | Ie.He | Ie.La | Ie.Md | Ie.Pe | Ie.Sa |  | Ie.Do | Ie.Cy | Ie.He | Ie.La | Ie.Md | Ie.Pe | Ie.Sa |
| Ie.Cy | 0.05994006 |  |  |  |  |  |  | Ie.Cy | 0.07132174 |  |  |  |  |  |  |
| Ie.He | **0.032967** | 0.1038961 |  |  |  |  |  | Ie.He | 0.25227614 | 0.20855545 |  |  |  |  |  |
| Ie.La | 0.21878122 | 0.74325674 | 0.136863137 |  |  |  |  | Ie.La | 0.05902363 | 0.03745724 | 0.20439403 |  |  |  |  |
| Ie.Md | 0.36263736 | 0.45554446 | **0.045954046** | 0.37062937 |  |  |  | Ie.Md | 0.06472334 | 0.05869873 | 0.25144317 | 0.06605893 |  |  |  |
| Ie.Pe | 0.47052947 | **0.002997** | **0.020979021** | **0.025974** | 0.21178821 |  |  | Ie.Pe | 0.04969331 | 0.10077055 | 0.27709736 | 0.08832166 | 0.07691534 |  |  |
| Ie.Sa | **0.000999** | **0.012987** | 0.126873127 | **0.007992** | **0.008991** | **0.000999** |  | Ie.Sa | 0.17966631 | 0.12597739 | 0.21142785 | 0.14285759 | 0.15035942 | 0.20364146 |  |
| Ie.Vi | 0.19180819 | **0.000999** | **0.00999001** | **0.001998** | 0.08791209 | 0.82017982 | **0.000999** | Ie.Vi | 0.05800421 | 0.11503468 | 0.29155241 | 0.10077488 | 0.08775433 | 0.03675285 | 0.21567857 |
| ***I. graellsii* (Posterior view)** | | |  |  |  |  |  |  |  |  |  |  |  |  |  |
| $p.value |  |  |  |  |  |  |  | $dist |  |  |  |  |  |  |  |
|  | Ig.Al | Ig.Ca | Ig.Ga | Ig.He | Ig.Ma | Ig.Mc | Ig.Pe |  | Ig.Al | Ig.Ca | Ig.Ga | Ig.He | Ig.Ma | Ig.Mc | Ig.Pe |
| Ig.Ca | 0.66533467 |  |  |  |  |  |  | Ig.Ca | 0.02582655 |  |  |  |  |  |  |
| Ig.Ga | 0.69230769 | 0.76523477 |  |  |  |  |  | Ig.Ga | 0.02690273 | 0.01844055 |  |  |  |  |  |
| Ig.He | 0.22577423 | 0.63436563 | 0.527472527 |  |  |  |  | Ig.He | 0.03651138 | 0.02005768 | 0.02348885 |  |  |  |  |
| Ig.Ma | 0.4995005 | 0.27572428 | 0.3996004 | **0.03996** |  |  |  | Ig.Ma | 0.02915428 | 0.02387652 | 0.02353394 | 0.03372036 |  |  |  |
| Ig.Mc | 0.31168831 | **0.035964** | **0.02997003** | **0.00999** | **0.03996** |  |  | Ig.Mc | 0.0352911 | 0.03511495 | 0.03955703 | 0.04183894 | 0.03576053 |  |  |
| Ig.Pe | 0.54845155 | 0.69430569 | 0.77022977 | 0.75224775 | 0.6003996 | 0.2947053 |  | Ig.Pe | 0.06321706 | 0.05073312 | 0.04771953 | 0.04915243 | 0.05570511 | 0.07741784 |  |
| Ig.Xu | 0.31968032 | **0.017982** | **0.002997003** | **0.000999** | **0.004995** | 0.32167832 | 0.20579421 | Ig.Xu | 0.0363507 | 0.04290118 | 0.04867273 | 0.05312637 | 0.04673618 | 0.03160133 | 0.08673353 |
| ***I. graellsii* (Lateral view)** | | |  |  |  |  |  |  |  |  |  |  |  |  |  |
| $p.value |  |  |  |  |  |  |  | $dist |  |  |  |  |  |  |  |
|  | Ig.Al | Ig.Ca | Ig.Ga | Ig.He | Ig.Ma | Ig.Mc | Ig.Pe |  | Ig.Al | Ig.Ca | Ig.Ga | Ig.He | Ig.Ma | Ig.Mc | Ig.Pe |
| Ig.Ca | 0.83616384 |  |  |  |  |  |  | Ig.Ca | 0.03452667 |  |  |  |  |  |  |
| Ig.Ga | 0.06793207 | **0.000999** |  |  |  |  |  | Ig.Ga | 0.07199564 | 0.08563742 |  |  |  |  |  |
| Ig.He | 0.58141858 | **0.043956** | 0.092907093 |  |  |  |  | Ig.He | 0.04367019 | 0.05416444 | 0.05124047 |  |  |  |  |
| Ig.Ma | 0.81018981 | 0.10889111 | **0.028971029** | 0.81718282 |  |  |  | Ig.Ma | 0.03574017 | 0.04415065 | 0.05972401 | 0.02792434 |  |  |  |
| Ig.Mc | 0.66633367 | **0.032967** | 0.124875125 | 0.22277722 | 0.48851149 |  |  | Ig.Mc | 0.04300638 | 0.05911595 | 0.05251837 | 0.04705921 | 0.03644202 |  |  |
| Ig.Pe | 0.97902098 | 0.73526474 | 0.925074925 | 0.96903097 | 0.96003996 | 0.97602398 |  | Ig.Pe | 0.05114012 | 0.0727287 | 0.05313253 | 0.04576843 | 0.04690375 | 0.04511751 |  |
| Ig.Xu | 0.50549451 | **0.041958** | 0.190809191 | 0.92907093 | 0.67632368 | 0.64735265 | 0.97502498 | Ig.Xu | 0.05039567 | 0.06484103 | 0.05206371 | 0.02898383 | 0.03544401 | 0.03831319 | 0.04551728 |

**Table S23**. Results of Procrustes ANOVA for centroid size of male caudal appendages of *Ischnura elegans* and *I. graellsii* between localities.

|  | Df | SS | MS | Rsq | F | Z | Pr(>F) |
| --- | --- | --- | --- | --- | --- | --- | --- |
| ***I. elegans* - Posterior view** |  |  |  |  |  |  |  |
| groups | 9 | 443615 | 49291 | 0.55628 | 25.352 | 8.3023 | **0.001** |
| Residuals | 182 | 353846 | 1944 | 0.44372 |  |  |  |
| Total | 191 | 797461 |  |  |  |  |  |
| ***I. elegans* - Lateral view** |  |  |  |  |  |  |  |
| groups | 9 | 235373 | 26152.6 | 0.41133 | 14.13 | 7.0401 | **0.001** |
| Residuals | 182 | 336848 | 1850.8 | 0.58867 |  |  |  |
| Total | 191 | 572221 |  |  |  |  |  |
| ***I. graellsii* - Posterior view** |  |  |  |  |  |  |  |
| groups | 6 | 79284 | 13214 | 0.24236 | 6.9309 | 4.8814 | **0.001** |
| Residuals | 130 | 247850 | 1906.5 | 0.75764 |  |  |  |
| Total | 136 | 327134 |  |  |  |  |  |
| ***I. graellsii* - Lateral view** |  |  |  |  |  |  |  |
| groups | 6 | 29617 | 4936.1 | 0.12458 | 3.0834 | 2.723 | **0.004** |
| Residuals | 130 | 208114 | 1600.9 | 0.87542 |  |  |  |
| Total | 136 | 237730 |  |  |  |  |  |

**Table S24**. Results of pairwise comparison of centroid size of male caudal appendages between *Ischnura elegans* and *I. graellsii* estimated between localities. Ie: *Ischnura elegans*, Ig: *I. graellsii*, Al: Alba, Ca: Cachadas, Cy: St. Cyprien, Do: Doniños, Ga: Gamillazo, He: Hervias, La: Laxe, Ma: Mateo, Md: Marais d’Orx, Pe: Perdiguero, Pl: Plaiaundi, Sa: Sabiere, Vi: Villar, Xu: Xuño.

|  | Posterior view | | | | | Lateral view | | | | |
| --- | --- | --- | --- | --- | --- | --- | --- | --- | --- | --- |
|  | d.obs | UCL (95%) | Zd | Pr(>d) | p.adjusted | d.obs | UCL (95%) | Zd | Pr(>d) | p.adjusted |
| ***I. elegans*** |  |  |  |  |  |  |  |  |  |  |
| Ie.Cy:Ie.Do | 3.913695 | 34.53757 | -1.06992 | 0.85 | 0.991023 | 3.4426562 | 29.7869 | -1.04676 | 0.842 | 0.890581 |
| Ie.Cy:Ie.He | 53.08702 | 100.9758 | 0.662393 | 0.275 | 0.479423 | 49.2117467 | 85.56209 | 0.73923 | 0.239 | 0.398333 |
| Ie.Cy:Ie.La | 51.44495 | 35.12327 | 2.33936 | **0.006** | **0.018** | 34.1169176 | 29.74128 | 1.867716 | **0.023** | 0.064688 |
| Ie.Cy:Ie.Md | 60.80588 | 35.8864 | 2.662313 | **0.004** | **0.01385** | 57.2885807 | 30.14837 | 2.907555 | **0.001** | **0.00409** |
| Ie.Cy:Ie.Pe | 50.54046 | 37.44875 | 2.116406 | **0.014** | **0.03706** | 41.7080101 | 31.82229 | 2.091393 | **0.013** | **0.04179** |
| Ie.Cy:Ie.Pl | 72.11519 | 40.1803 | 2.698512 | **0.002** | **0.0075** | 74.8468285 | 34.93539 | 3.107107 | **0.001** | **0.00409** |
| Ie.Cy:Ie.Sa | 51.31104 | 34.76965 | 2.379299 | **0.002** | **0.0075** | 49.9091736 | 28.48823 | 2.702853 | **0.001** | **0.00409** |
| Ie.Cy:Ie.Vi | 161.4691 | 36.17769 | 5.175539 | **0.001** | **0.00563** | 113.0419386 | 29.53941 | 4.996196 | **0.001** | **0.00409** |
| Ie.Cy:Ie.Xu | 9.923019 | 75.40579 | -0.84101 | 0.783 | 0.952297 | 6.9677559 | 62.87146 | -1.00776 | 0.843 | 0.890581 |
| Ie.Do:Ie.He | 49.17332 | 101.6332 | 0.560155 | 0.298 | 0.487241 | 45.7690905 | 85.1016 | 0.627767 | 0.274 | 0.425172 |
| Ie.Do:Ie.La | 47.53126 | 36.50011 | 2.112747 | **0.006** | **0.018** | 30.6742613 | 29.34589 | 1.641495 | **0.043** | 0.09675 |
| Ie.Do:Ie.Md | 56.89218 | 38.52734 | 2.374322 | **0.002** | **0.0075** | 53.8459245 | 32.08126 | 2.750919 | **0.001** | **0.00409** |
| Ie.Do:Ie.Pe | 46.62677 | 39.83295 | 1.897881 | **0.022** | **0.055** | 38.2653539 | 33.61376 | 1.906659 | **0.027** | 0.071471 |
| Ie.Do:Ie.Pl | 68.20149 | 42.73193 | 2.448133 | **0.002** | **0.0075** | 71.4041722 | 34.56931 | 3.14857 | **0.001** | **0.00409** |
| Ie.Do:Ie.Sa | 47.39734 | 34.38176 | 2.16474 | **0.007** | **0.01969** | 46.4665173 | 30.08428 | 2.472282 | **0.004** | 0.013846 |
| Ie.Do:Ie.Vi | 157.5554 | 35.04294 | 5.157765 | **0.001** | **0.00563** | 109.5992824 | 31.11041 | 4.436659 | **0.001** | **0.00409** |
| Ie.Do:Ie.Xu | 13.83671 | 73.24014 | -0.58544 | 0.719 | 0.89875 | 3.5250997 | 61.7836 | -1.45207 | 0.915 | 0.935795 |
| Ie.He:Ie.La | 1.642062 | 94.10008 | -1.84606 | 0.965 | 0.991023 | 15.0948292 | 82.38903 | -0.57986 | 0.708 | 0.816923 |
| Ie.He:Ie.Md | 7.718863 | 94.02675 | -1.18407 | 0.879 | 0.991023 | 8.076834 | 81.95626 | -1.07597 | 0.84 | 0.890581 |
| Ie.He:Ie.Pe | 2.546556 | 97.38076 | -1.75832 | 0.951 | 0.991023 | 7.5037366 | 82.23172 | -1.14224 | 0.851 | 0.890581 |
| Ie.He:Ie.Pl | 19.02817 | 102.3269 | -0.43846 | 0.675 | 0.867857 | 25.6350817 | 88.56298 | -0.09086 | 0.552 | 0.709714 |
| Ie.He:Ie.Sa | 1.775977 | 97.29204 | -1.90951 | 0.969 | 0.991023 | 0.6974268 | 82.9845 | -2.12782 | 0.986 | 0.986 |
| Ie.He:Ie.Vi | 108.3821 | 97.84235 | 1.71173 | **0.035** | 0.07875 | 63.8301919 | 83.26589 | 1.130412 | 0.123 | 0.240652 |
| Ie.He:Ie.Xu | 63.01003 | 118.4572 | 0.634986 | 0.277 | 0.479423 | 42.2439909 | 103.0569 | 0.340394 | 0.378 | 0.500294 |
| Ie.La:Ie.Md | 9.360925 | 36.52827 | -0.25403 | 0.601 | 0.836471 | 23.1716632 | 29.7065 | 1.13476 | 0.141 | 0.264375 |
| Ie.La:Ie.Pe | 0.904494 | 38.91048 | -1.86757 | 0.965 | 0.991023 | 7.5910926 | 32.9804 | -0.38481 | 0.642 | 0.760263 |
| Ie.La:Ie.Pl | 20.67023 | 41.36188 | 0.557979 | 0.314 | 0.487241 | 40.7299109 | 34.14443 | 1.907706 | **0.018** | 0.054 |
| Ie.La:Ie.Sa | 0.133916 | 33.47987 | -2.32914 | 0.995 | 0.995 | 15.792256 | 28.69797 | 0.639218 | 0.287 | 0.4305 |
| Ie.La:Ie.Vi | 110.0241 | 34.67607 | 4.181063 | **0.001** | **0.00563** | 78.9250211 | 29.29712 | 3.847974 | **0.001** | **0.00409** |
| Ie.La:Ie.Xu | 61.36797 | 72.91958 | 1.27582 | 0.095 | 0.194318 | 27.1491617 | 62.49423 | 0.344924 | 0.374 | 0.500294 |
| Ie.Md:Ie.Pe | 10.26542 | 40.59949 | -0.3409 | 0.632 | 0.836471 | 15.5805706 | 34.4843 | 0.424573 | 0.359 | 0.500294 |
| Ie.Md:Ie.Pl | 11.30931 | 39.9058 | -0.24736 | 0.583 | 0.836471 | 17.5582478 | 33.04324 | 0.556563 | 0.315 | 0.457258 |
| Ie.Md:Ie.Sa | 9.49484 | 36.97208 | -0.29504 | 0.627 | 0.836471 | 7.3794072 | 29.79798 | -0.31431 | 0.632 | 0.760263 |
| Ie.Md:Ie.Vi | 100.6632 | 35.96894 | 3.956028 | **0.001** | **0.00563** | 55.7533579 | 30.28593 | 2.931508 | **0.001** | **0.00409** |
| Ie.Md:Ie.Xu | 70.7289 | 76.48887 | 1.490072 | 0.064 | 0.137143 | 50.3208248 | 62.29701 | 1.192242 | 0.118 | 0.240652 |
| Ie.Pe:Ie.Pl | 21.57473 | 43.03914 | 0.550861 | 0.327 | 0.4905 | 33.1388184 | 35.70095 | 1.398347 | 0.083 | 0.177857 |
| Ie.Pe:Ie.Sa | 0.770579 | 37.45975 | -1.87968 | 0.967 | 0.991023 | 8.2011634 | 32.85537 | -0.29099 | 0.626 | 0.760263 |
| Ie.Pe:Ie.Vi | 110.9286 | 38.30551 | 3.790222 | **0.001** | **0.00563** | 71.3339285 | 33.48513 | 3.000794 | **0.001** | **0.00409** |
| Ie.Pe:Ie.Xu | 60.46348 | 79.62514 | 1.176669 | 0.123 | 0.230625 | 34.7402542 | 64.70819 | 0.636016 | 0.266 | 0.425172 |
| Ie.Pl:Ie.Sa | 20.80415 | 40.66854 | 0.539984 | 0.312 | 0.487241 | 24.9376549 | 35.99264 | 1.005207 | 0.169 | 0.3042 |
| Ie.Pl:Ie.Vi | 89.35389 | 40.16592 | 3.329796 | **0.001** | **0.00563** | 38.1951102 | 34.90156 | 1.747842 | **0.034** | 0.085 |
| Ie.Pl:Ie.Xu | 82.03821 | 76.60369 | 1.664502 | **0.035** | 0.07875 | 67.8790726 | 63.72408 | 1.645811 | **0.042** | 0.09675 |
| Ie.Sa:Ie.Vi | 110.158 | 35.67902 | 4.115121 | **0.001** | **0.00563** | 63.1327651 | 28.88871 | 3.196939 | **0.001** | **0.00409** |
| Ie.Sa:Ie.Xu | 61.23406 | 75.14058 | 1.267614 | 0.105 | 0.205435 | 42.9414177 | 64.3279 | 0.976215 | 0.181 | 0.313269 |
| Ie.Vi:Ie.Xu | 171.3921 | 73.51605 | 3.258901 | **0.001** | **0.00563** | 106.0741827 | 63.21071 | 2.604618 | **0.004** | **0.01385** |
| ***I. graellsii*** |  |  |  |  |  |  |  |  |  |  |
| Ig.Al:Ig.Ca | 10.61125 | 29.80074 | 0.105286 | 0.469 | 0.518368 | 45.5601298 | 23.73152 | 2.891826 | **0.001** | **0.021** |
| Ig.Al:Ig.Ga | 32.82556 | 34.73929 | 1.44541 | 0.075 | 0.121154 | 19.816769 | 28.93845 | 0.949825 | 0.18 | 0.290769 |
| Ig.Al:Ig.He | 61.8037 | 29.83625 | 3.059923 | **0.001** | **0.00525** | 27.0872247 | 24.2469 | 1.775476 | **0.031** | 0.1302 |
| Ig.Al:Ig.Ma | 58.82841 | 32.04033 | 2.830142 | **0.001** | **0.00525** | 33.5022766 | 25.48531 | 2.131729 | **0.012** | 0.11025 |
| Ig.Al:Ig.Pl | 130.5586 | 98.58583 | 2.226819 | **0.006** | **0.0252** | 94.7059942 | 82.77176 | 1.88938 | **0.021** | 0.11025 |
| Ig.Al:Ig.Xu | 34.42051 | 31.4089 | 1.794538 | **0.033** | 0.099 | 20.4245411 | 25.32757 | 1.215778 | 0.116 | 0.221455 |
| Ig.Ca:Ig.Ga | 22.21431 | 29.51541 | 1.055092 | 0.153 | 0.191333 | 25.7433608 | 26.06201 | 1.546327 | 0.054 | 0.179667 |
| Ig.Ca:Ig.He | 51.19246 | 24.74834 | 3.241189 | **0.001** | **0.00525** | 18.4729051 | 19.99586 | 1.440025 | 0.074 | 0.179667 |
| Ig.Ca:Ig.Ma | 48.21717 | 26.48575 | 2.822149 | **0.001** | **0.00525** | 12.0578532 | 21.56587 | 0.655215 | 0.269 | 0.36225 |
| Ig.Ca:Ig.Pl | 119.9473 | 95.77851 | 2.06999 | **0.012** | **0.042** | 49.1458644 | 82.46049 | 0.774908 | 0.214 | 0.321 |
| Ig.Ca:Ig.Xu | 23.80926 | 25.07074 | 1.512859 | 0.066 | 0.1155 | 25.1355887 | 20.76908 | 1.935598 | **0.017** | 0.11025 |
| Ig.Ga:Ig.He | 28.97815 | 30.79561 | 1.476694 | 0.066 | 0.1155 | 7.2704557 | 26.55854 | -0.25069 | 0.603 | 0.63315 |
| Ig.Ga:Ig.Ma | 26.00285 | 33.29273 | 1.185758 | 0.122 | 0.1708 | 13.6855076 | 26.88289 | 0.469771 | 0.352 | 0.434824 |
| Ig.Ga:Ig.Pl | 97.733 | 98.37067 | 1.577116 | 0.053 | 0.1134 | 74.8892252 | 82.70983 | 1.453643 | 0.077 | 0.179667 |
| Ig.Ga:Ig.Xu | 1.594949 | 31.62502 | -1.52382 | 0.933 | 0.933 | 0.6077721 | 27.34881 | -1.81735 | 0.958 | 0.958 |
| Ig.He:Ig.Ma | 2.975291 | 26.63456 | -1.01235 | 0.831 | 0.87255 | 6.4150519 | 22.12266 | -0.11742 | 0.552 | 0.610105 |
| Ig.He:Ig.Pl | 68.75485 | 96.99121 | 0.978067 | 0.164 | 0.191333 | 67.6187695 | 79.4753 | 1.30143 | 0.102 | 0.2142 |
| Ig.He:Ig.Xu | 27.3832 | 26.27344 | 1.666074 | **0.043** | 0.112875 | 6.6626836 | 21.35301 | -0.10967 | 0.539 | 0.610105 |
| Ig.Ma:Ig.Pl | 71.73015 | 97.91702 | 1.034875 | 0.16 | 0.191333 | 61.2037176 | 82.8727 | 1.098764 | 0.132 | 0.231 |
| Ig.Ma:Ig.Xu | 24.40791 | 28.25925 | 1.361727 | 0.092 | 0.138 | 13.0777355 | 23.84792 | 0.664244 | 0.276 | 0.36225 |
| Ig.Pl:Ig.Xu | 96.13805 | 96.79749 | 1.580025 | 0.054 | 0.1134 | 74.2814531 | 82.55741 | 1.46258 | 0.075 | 0.179667 |

**Table S25**. Results of Procrustes ANOVA for centroid size of female prothorax of *Ischnura elegans* and *I. graellsii* between localities.

|  | Df | SS | MS | Rsq | F | Z | Pr(>F) |
| --- | --- | --- | --- | --- | --- | --- | --- |
| ***I. elegans* - Posterior view** |  |  |  |  |  |  |  |
| groups | 7 | 115433 | 16490.4 | 0.46036 | 15.355 | 7.325 | **0.001** |
| Residuals | 126 | 135315 | 1073.9 | 0.53964 |  |  |  |
| Total | 133 | 250748 |  |  |  |  |  |
| ***I. elegans* - Lateral view** |  |  |  |  |  |  |  |
| groups | 7 | 79379 | 11339.9 | 0.47765 | 16.46 | 8.6434 | **0.001** |
| Residuals | 126 | 86807 | 688.9 | 0.52235 |  |  |  |
| Total | 133 | 166186 |  |  |  |  |  |
| ***I. graellsii* - Posterior view** |  |  |  |  |  |  |  |
| groups | 7 | 10625 | 1517.9 | 0.04891 | 0.933 | 0.068263 | 0.469 |
| Residuals | 127 | 206625 | 1627 | 0.95109 |  |  |  |
| Total | 134 | 217251 |  |  |  |  |  |
| ***I. graellsii* - Lateral view** |  |  |  |  |  |  |  |
| groups | 7 | 5267 | 752.45 | 0.10872 | 2.213 | 1.6594 | **0.044** |
| Residuals | 127 | 43181 | 340.01 | 0.89128 |  |  |  |
| Total | 134 | 48449 |  |  |  |  |  |

**Table S26**. Results of pairwise comparison of centroid size of female prothorax of *Ischnura elegans* and *I. graellsii* estimated between localities. Ie: *Ischnura elegans*, Ig: *I. graellsii*, Al: Alba, Ca: Cachadas, Cy: St. Cyprien, Do: Doniños, Ga: Gamillazo, He: Hervias, La: Laxe, Ma: Mateo, Mc: Saïdia, Pe: Perdiguero, Sa: Sabiere, Vi: Villar, Xu: Xuño.

|  | Posterior view | | | | | Lateral view | | | | |
| --- | --- | --- | --- | --- | --- | --- | --- | --- | --- | --- |
|  | d.obs | UCL (95%) | Zd | Pr(>d) | p.adjusted | d.obs | UCL (95%) | Zd | Pr(>d) | p.adjusted |
| ***I. elegans*** |  |  |  |  |  |  |  |  |  |  |
| Ie.Do:Ie.Fr | 28.63611 | 24.45689 | 1.944348 | **0.018** | 0.056 | 12.225906 | 19.75724 | 0.795961 | 0.239 | 0.446133 |
| Ie.Do:Ie.He | 13.26864 | 83.36821 | -0.84494 | 0.792 | 0.852923 | 33.069901 | 68.71605 | 0.234508 | 0.43 | 0.602 |
| Ie.Do:Ie.La | 20.00947 | 24.77489 | 1.258776 | 0.105 | 0.21 | 5.553883 | 20.03938 | -0.16285 | 0.57 | 0.719478 |
| Ie.Do:Ie.Md | 48.55451 | 32.81257 | 2.321953 | **0.004** | **0.014** | 2.031132 | 26.39206 | -1.22377 | 0.879 | 0.879 |
| Ie.Do:Ie.Pe | 72.92625 | 25.83418 | 4.08544 | **0.001** | **0.00467** | 46.52113 | 20.85005 | 3.454526 | **0.001** | **0.004** |
| Ie.Do:Ie.Sa | 52.25446 | 35.68697 | 2.37318 | **0.004** | **0.014** | 11.441682 | 27.75756 | 0.170056 | 0.458 | 0.610667 |
| Ie.Do:Ie.Vi | 82.01446 | 23.73058 | 4.383015 | **0.001** | **0.00467** | 49.763132 | 19.86162 | 3.552794 | **0.001** | **0.004** |
| Ie.Fr:Ie.He | 15.36747 | 85.38205 | -0.67501 | 0.749 | 0.83888 | 20.843995 | 67.57307 | -0.34833 | 0.627 | 0.7315 |
| Ie.Fr:Ie.La | 8.62664 | 23.51059 | 0.1423 | 0.455 | 0.553913 | 17.779789 | 20.17608 | 1.410752 | 0.079 | 0.170154 |
| Ie.Fr:Ie.Md | 19.9184 | 33.17681 | 0.780315 | 0.238 | 0.350737 | 14.257038 | 26.66671 | 0.596387 | 0.29 | 0.484235 |
| Ie.Fr:Ie.Pe | 44.29014 | 26.14437 | 2.601485 | **0.001** | **0.00467** | 58.747037 | 21.01507 | 3.94976 | **0.001** | **0.004** |
| Ie.Fr:Ie.Sa | 23.61835 | 34.49905 | 0.968582 | 0.191 | 0.314588 | 23.667588 | 28.67131 | 1.259681 | 0.113 | 0.226 |
| Ie.Fr:Ie.Vi | 53.37835 | 24.90197 | 3.172444 | **0.001** | **0.00467** | 61.989039 | 20.20221 | 4.217094 | **0.001** | **0.004** |
| Ie.He:Ie.La | 6.740828 | 85.69714 | -1.28867 | 0.886 | 0.886 | 38.623783 | 69.67548 | 0.479782 | 0.359 | 0.558444 |
| Ie.He:Ie.Md | 35.28586 | 86.35238 | 0.163019 | 0.45 | 0.553913 | 35.101033 | 70.32391 | 0.31687 | 0.415 | 0.602 |
| Ie.He:Ie.Pe | 59.65761 | 86.162 | 0.928522 | 0.205 | 0.318889 | 79.591031 | 70.15493 | 1.842343 | **0.023** | 0.053667 |
| Ie.He:Ie.Sa | 38.98582 | 91.33955 | 0.298626 | 0.401 | 0.534667 | 44.511583 | 72.61112 | 0.651145 | 0.294 | 0.484235 |
| Ie.He:Ie.Vi | 68.74582 | 84.85225 | 1.163323 | 0.15 | 0.2625 | 82.833033 | 68.89196 | 1.957402 | **0.019** | **0.04836** |
| Ie.La:Ie.Md | 28.54504 | 31.87092 | 1.382287 | 0.077 | 0.179667 | 3.52275 | 26.79829 | -0.85684 | 0.79 | 0.819259 |
| Ie.La:Ie.Pe | 52.91678 | 27.2271 | 3.006024 | **0.001** | **0.00467** | 40.967248 | 21.62659 | 3.019128 | **0.001** | **0.004** |
| Ie.La:Ie.Sa | 32.24499 | 34.43868 | 1.480955 | 0.068 | 0.173091 | 5.887799 | 27.87629 | -0.57551 | 0.702 | 0.78624 |
| Ie.La:Ie.Vi | 62.00499 | 24.69656 | 3.598525 | **0.001** | **0.00467** | 44.20925 | 20.42318 | 3.238051 | **0.001** | **0.004** |
| Ie.Md:Ie.Pe | 24.37175 | 34.18461 | 1.085278 | 0.147 | 0.2625 | 44.489998 | 26.45359 | 2.612783 | **0.003** | **0.0105** |
| Ie.Md:Ie.Sa | 3.699956 | 39.77697 | -1.15067 | 0.853 | 0.884593 | 9.41055 | 32.14214 | -0.18711 | 0.591 | 0.719478 |
| Ie.Md:Ie.Vi | 33.45996 | 32.34556 | 1.660943 | **0.043** | 0.1204 | 47.732 | 26.15169 | 2.859097 | **0.001** | **0.004** |
| Ie.Pe:Ie.Sa | 20.67179 | 35.02843 | 0.681623 | 0.271 | 0.3794 | 35.079448 | 29.47151 | 1.939931 | **0.013** | **0.0364** |
| Ie.Pe:Ie.Vi | 9.088207 | 25.16728 | 0.057676 | 0.509 | 0.593833 | 3.242002 | 20.39924 | -0.72901 | 0.75 | 0.807692 |
| Ie.Sa:Ie.Vi | 29.76 | 34.47945 | 1.321783 | 0.088 | 0.189538 | 38.32145 | 27.93085 | 2.170629 | **0.006** | **0.01867** |
| ***I. graellsii*** |  |  |  |  |  |  |  |  |  |  |
| Ig.Al:Ig.Ca | 13.8203 | 28.04158 | 0.458012 | 0.347 | 0.693 | 8.3403434 | 14.46682 | 0.730537 | 0.25 | 0.499059 |
| Ig.Al:Ig.Ga | 16.33477 | 31.63578 | 0.553237 | 0.309 | 0.693 | 0.9450524 | 15.71428 | -1.42035 | 0.914 | 0.914 |
| Ig.Al:Ig.He | 25.15633 | 30.92602 | 1.179713 | 0.123 | 0.693 | 11.7081608 | 15.5257 | 1.167085 | 0.121 | 0.4235 |
| Ig.Al:Ig.Ma | 23.03909 | 30.3068 | 1.141536 | 0.126 | 0.693 | 12.3228373 | 14.03735 | 1.311304 | 0.106 | 0.4235 |
| Ig.Al:Ig.Mc | 0.162263 | 34.5214 | -2.28549 | 0.989 | 0.989 | 5.3822665 | 16.40844 | 0.00993 | 0.494 | 0.728 |
| Ig.Al:Ig.Pe | 48.17356 | 82.52283 | 0.794145 | 0.246 | 0.693 | 9.1157061 | 40.00398 | -0.24189 | 0.597 | 0.759818 |
| Ig.Al:Ig.Xu | 17.77027 | 33.81959 | 0.579254 | 0.298 | 0.693 | 10.297087 | 15.50727 | 0.883186 | 0.189 | 0.481091 |
| Ig.Ca:Ig.Ga | 2.514466 | 23.07204 | -1.03962 | 0.842 | 0.926154 | 7.395291 | 10.85456 | 0.933224 | 0.184 | 0.481091 |
| Ig.Ca:Ig.He | 11.33603 | 22.95336 | 0.5009 | 0.316 | 0.693 | 3.3678174 | 11.1473 | -0.13834 | 0.56 | 0.746667 |
| Ig.Ca:Ig.Ma | 9.218785 | 21.15791 | 0.335031 | 0.396 | 0.693 | 3.9824939 | 10.61977 | 0.172366 | 0.466 | 0.724889 |
| Ig.Ca:Ig.Mc | 13.65804 | 25.46603 | 0.605123 | 0.285 | 0.693 | 13.72261 | 11.96084 | 1.867998 | **0.021** | 0.1568 |
| Ig.Ca:Ig.Pe | 34.35326 | 77.6813 | 0.394686 | 0.356 | 0.693 | 17.4560496 | 39.83442 | 0.519707 | 0.296 | 0.499059 |
| Ig.Ca:Ig.Xu | 3.949964 | 26.94671 | -0.76024 | 0.76 | 0.886667 | 1.9567436 | 12.51559 | -0.67356 | 0.739 | 0.865667 |
| Ig.Ga:Ig.He | 8.821561 | 25.05297 | 0.073505 | 0.499 | 0.735368 | 10.7631084 | 11.9621 | 1.445662 | 0.081 | 0.378 |
| Ig.Ga:Ig.Ma | 6.704319 | 22.54629 | -0.16083 | 0.568 | 0.757333 | 11.3777849 | 10.30256 | 1.74931 | **0.028** | 0.1568 |
| Ig.Ga:Ig.Mc | 16.17251 | 27.2628 | 0.747677 | 0.245 | 0.693 | 6.327319 | 12.15608 | 0.523661 | 0.303 | 0.499059 |
| Ig.Ga:Ig.Pe | 31.83879 | 76.92241 | 0.303388 | 0.392 | 0.693 | 10.0607586 | 40.64856 | -0.08235 | 0.545 | 0.746667 |
| Ig.Ga:Ig.Xu | 1.435498 | 28.7964 | -1.48097 | 0.92 | 0.954074 | 9.3520346 | 12.33824 | 1.053621 | 0.151 | 0.469778 |
| Ig.He:Ig.Ma | 2.117242 | 22.95289 | -1.08906 | 0.86 | 0.926154 | 0.6146765 | 11.02164 | -1.33116 | 0.889 | 0.914 |
| Ig.He:Ig.Mc | 24.99407 | 26.28538 | 1.500216 | 0.065 | 0.693 | 17.0904273 | 12.34182 | 2.202543 | **0.009** | 0.126 |
| Ig.He:Ig.Pe | 23.01723 | 80.72692 | -0.13329 | 0.552 | 0.757333 | 20.8238669 | 39.94745 | 0.716993 | 0.243 | 0.499059 |
| Ig.He:Ig.Xu | 7.386062 | 27.97483 | -0.25421 | 0.607 | 0.772545 | 1.4110738 | 12.79592 | -1.06002 | 0.843 | 0.907846 |
| Ig.Ma:Ig.Mc | 22.87682 | 25.31444 | 1.429122 | 0.073 | 0.693 | 17.7051038 | 11.97652 | 2.348079 | **0.004** | 0.112 |
| Ig.Ma:Ig.Pe | 25.13447 | 79.52933 | 0.024162 | 0.472 | 0.734222 | 21.4385434 | 40.97151 | 0.747904 | 0.217 | 0.499059 |
| Ig.Ma:Ig.Xu | 5.268821 | 26.56641 | -0.58317 | 0.715 | 0.870435 | 2.0257503 | 12.67569 | -0.72019 | 0.742 | 0.865667 |
| Ig.Mc:Ig.Pe | 48.0113 | 80.53857 | 0.773644 | 0.251 | 0.693 | 3.7334396 | 40.71 | -0.94531 | 0.828 | 0.907846 |
| Ig.Mc:Ig.Xu | 17.608 | 30.62688 | 0.743959 | 0.246 | 0.693 | 15.6793535 | 13.6909 | 1.905104 | **0.025** | 0.1568 |
| Ig.Pe:Ig.Xu | 30.40329 | 79.35149 | 0.196276 | 0.433 | 0.713176 | 19.4127931 | 39.89171 | 0.586195 | 0.287 | 0.499059 |

**Table S27**. Relationship between the CS and the hindwing length of males and females of *I. elegans* and *I. graellsii*.

|  | Locality | R pearson | *p* pearson |
| --- | --- | --- | --- |
| Males | Cachadas | 0.84948569 | **0.0018647** |
|  | Cyprien | 0.34925232 | 0.32257773 |
|  | Gamillazo | -0.06802949 | 0.8518723 |
|  | Hervias | 0.63802805 | **0.0471487** |
|  | Laxe | 0.29231425 | 0.41245674 |
|  | Perdiguero | -0.45935043 | 0.1816989 |
| Females | Cachadas | 0.23789128 | 0.50807621 |
|  | Cyprien | 0.3481747 | 0.3241754 |
|  | Gamillazo | 0.2857642 | 0.42348545 |
|  | Hervias | -0.36783584 | 0.295676 |
|  | Laxe | 0.65615145 | **0.0393603** |
|  | Perdiguero | 0.75512086 | **0.0115643** |

**Table S28**. Proportion of the relationship between male and female genitalia of *I. elegans* and *I. graellsii* within each zone studied.

|  |  | CS | Shape |  | CS | Shape |
| --- | --- | --- | --- | --- | --- | --- |
| Posterior view | I.e_Alo | 19% | 3% | I.g_Alo | 19% | 7% |
|  | I.e_NW | 13% | 6% | I.g_NW | 6% | 3% |
|  | I.e_NCM | 7% | 6% | I.g_NCM | 1% | 5% |
| Lateral view | I.e_Alo | 20% | 5% | I.g_Alo | 4% | 5% |
|  | I.e_NW | 2% | 10% | I.g_NW | 5% | 8% |
|  | I.e_NCM | 2% | 4% | I.g_NCM | 6% | 5% |
